# NEKL-4 regulates microtubule stability and mitochondrial health in *C. elegans* ciliated neurons

**DOI:** 10.1101/2024.02.14.580304

**Authors:** Kaiden M. Power, Ken C. Nguyen, Andriele Silva, Shaneen Singh, David H. Hall, Christopher Rongo, Maureen M. Barr

## Abstract

Ciliopathies are often caused by defects in the ciliary microtubule core. Glutamylation is abundant in cilia, and its dysregulation may contribute to ciliopathies and neurodegeneration. Mutation of the deglutamylase CCP1 causes infantile-onset neurodegeneration. In *C. elegans, ccpp-1* loss causes age-related ciliary degradation that is suppressed by mutation in the conserved NEK10 homolog *nekl-4*. NEKL-4 is absent from cilia, yet negatively regulates ciliary stability via an unknown, glutamylation-independent mechanism. We show that NEKL-4 was mitochondria-associated. *nekl-4* mutants had longer mitochondria, a higher baseline mitochondrial oxidation state, and suppressed *ccpp-1* mutant lifespan extension in response to oxidative stress. A kinase-dead *nekl-4(KD)* mutant ectopically localized to *ccpp-1* cilia and rescued degenerating microtubule doublet B-tubules. A nondegradable *nekl-4(PESTΔ)* mutant resembled the *ccpp-1* mutant with dye filling defects and B-tubule breaks. The *nekl-4(PESTΔ)* Dyf phenotype was suppressed by mutation in the depolymerizing kinesin-8 KLP-13/KIF19A. We conclude that NEKL-4 influences ciliary stability by activating ciliary kinesins and promoting mitochondrial homeostasis.

**Summary:** Neurodegeneration and ciliary degeneration are caused by mutation of the deglutamylase CCP1/CCPP-1 in humans and *C. elegans*. The conserved NIMA-related kinase NEKL-4/NEK10 can suppress or promote degeneration in an activity-dependent manner that involves cilia-mitochondria communication and that is independent of glutamylation.

## Introduction

Cilia are microtubule-based organelles that project from most cell types in the body, including those in the nervous system (Lee and Gleeson, 2011). In the brain, motile cilia generate cerebrospinal flow (Spassky and Meunier, 2017) and neuronal primary cilia form synapses (Sheu et al., 2022). Ciliopathies, disorders of cilia, comprise a diverse group of diseases ranging from single-organ dysfunction (eg. polycystic kidney disease, retinitis pigmentosa) to congenital malformations and developmental defects that affect multiple organ systems including the brain (eg. Joubert syndrome, Bardet–Biedl syndrome) (Mill et al., 2023). The causes of ciliopathies vary widely, but many involve ultrastructural abnormalities in the microtubule-based structural core of the cilium, called the axoneme. The axoneme is composed of a ring of nine outer A-B doublet microtubules surrounding a pair of central singlets or no central pair - the “9 + 2” or “9 + 0” formation in motile or primary/sensory cilia, respectively (Mill et al., 2023).

Ciliary microtubule structure and microtubule-based transport are regulated by the “Tubulin Code” via tubulin <u>p</u>ost-translational <u>m</u>odifications (PTMs) that are added and removed by enzymatic “writers’’ and “erasers” (McKenna et al., 2023). Glutamylation is an evolutionarily conserved PTM that occurs in cilia from protozoa to mammals. Dysregulated glutamylation (hypo- or hyperglutamylation) leads to neurodegeneration and ciliopathies such as idiopathic scoliosis (Mathieu et al., 2021), retinitis pigmentosa (Cehajic-Kapetanovic et al., 2022), and Joubert syndrome (He et al., 2018). Tubulin PTMs regulate microtubule-based transport and are interpreted by “readers” that include kinesin motors. The localization of proteins and other molecular cargoes along microtubules is of vital importance in the long processes of neurons, and dysregulation of this process may contribute to neurodegeneration (Guo et al., 2020).

Mitochondria are transported along microtubules, making microtubule-based transport essential since mitochondrial ATP is required at synapses that may be a great distance from the neuronal cell body (Panchal and Tiwari, 2019; Cheng and Sheng, 2021). Hyperglutamylation caused by mutation of the deglutamylase CCP1 results in early-onset neurodegeneration and defects in microtubule-based mitochondrial transport (Shashi et al., 2018; Magiera et al., 2018; Gilmore-Hall et al., 2019; Li et al., 2020).

The *C. elegans* CCP1 homolog *ccpp-1* regulates microtubule stability in sensory cilia (O’Hagan et al., 2011). In the cilia of *ccpp-1* mutants, B-tubules degenerate in amphid chemosensory neurons in an age-dependent and progressive manner. Glutamylation occurs preferentially on the B-tubule (Lechtreck and Geimer, 2000; Kubo et al., 2010; Kubo and Oda, 2017; Orbach and Howard, 2019; Alvarez Viar et al., 2023), consistent with *ccpp-1* hyperglutamylation-induced B-tubule defects. Previously, we identified the NIMA-related kinase NEK10/NEKL-4 as a genetic interactor of the CCPP-1 deglutamylase (Power et al., 2020). Loss-of-function mutations of *nekl-4* suppress *ccpp-1* progressive ciliary degeneration, as measured by fluorescent lipophilic <u>dy</u>e filling defects (Dyf) of the ciliated neurons. Conversely, overexpression of NEKL-4 results in a progressive Dyf phenotype (Power et al., 2020; Li et al., 2021a). Endogenous NEKL-4 protein does not localize to the cilium or influence ciliary tubulin glutamylation, and the NEKL-4 mechanism of action remained unclear (Power et al., 2020). The mammalian NEKL-4 ortholog NEK10 has potential roles in cilia motility in airway epithelial cilia (Chivukula et al., 2020; Al Mutairi et al., 2020) and regulating mitochondria localization and homeostasis in mammalian cell culture (Peres de Oliveira et al., 2020), but how these processes are connected remains elusive.

To understand NEKL-4/NEK10 function and to determine where and how NEKL-4 regulates ciliary stability, we used CRISPR/Cas9 genomic editing to generate NEKL-4/NEK10 endogenously tagged fluorescent reporters and targeted mutations. We engineered individual mutations in the active site or the PEST domain of the NEKL-4 endogenously tagged fluorescent reporter strain to generate predicted kinase dead NEKL-4(KD) and non-degradable NEKL-4(PESTΔ) mutants. We discovered that NEKL-4 was mitochondria-associated, and that *nekl-4* mutants displayed in longer mitochondria, a higher baseline mitochondrial oxidation state, and suppression of the *ccpp-1* mutant lifespan extension in response to oxidative stress. NEKL-4(KD) also localized to mitochondria, but ectopically localized to cilia of *ccpp-1* deglutamylase and *ttll-11* glutamylase mutants. Additionally, *nekl-4(KD)* suppressed the B-tubule degradation of *ccpp-1* cilia and prevented ciliary degeneration. Conversely, the *nekl-4(PESTΔ)* mutant resembled the *ccpp-1* mutant with dye filling defects and ultrastructural defects including B-tubule breaks. The *nekl-4(PESTΔ)* Dyf defect was suppressed by mutation of the depolymerizing kinesin-8 *klp-13/KIF19A*, suggesting that NEKL-4 may regulate ciliary stability by regulating this “reader” of the Tubulin Code. Combined, our data indicate that NEKL-4 negatively influences ciliary stability by two distinct but related mechanisms: activation of ciliary kinesins and reduction of mitochondrial stress.

## Results

### Kinase-dead NEKL-4(KD) suppresses the ccpp-1Δ Dye-filling phenotype and ectopically localizes to cilia in ccpp-1Δ and ttll-11Δ mutants

Our previous study was performed with null *nekl-4* alleles that were either an early stop codon (*my31*) or a partial deletion that eliminates the ATP binding site in the kinase domain (*tm4910*, referred to as *nekl-4Δ*) (Power et al., 2020). We wanted to determine if loss of NEKL-4 kinase activity was required or sufficient for suppression of the *ccpp-1Δ* Dyf phenotype. To accomplish this, we used CRISPR/Cas9 to engineer a kinase-dead *nekl-4* allele by mutating the active site predicted by InterPro from D591 to A591 (**Fig 1a**). Asp to Ala substitutions have also been used in other studies to create kinase-dead proteins (Lew et al., 2009; Muñoz et al., 2018). This active site-disrupted allele will be referred to as *nekl-4(KD)* and is also endogenously tagged with mNeonGreen. This allele was used for most experiments other than those in which the fluorescent tag would interfere. *In silico* analysis based on the mammalian homolog NEK10 indicated that the D591A substitution does not alter the overall structure of NEKL-4 outside of the active site (**Fig 1b, Fig S1**). We performed a dye-filling assay to assess if *nekl-4(KD)* suppressed the *ccpp-1* dye-filling phenotype. We found that *nekl-4(KD)* suppressed *ccpp-1Δ* Dyf to a similar extent as *nekl-4(my31)* and *nekl-4Δ* (**Fig 1c-d**; (Power et al., 2020)).

**Figure 1.**
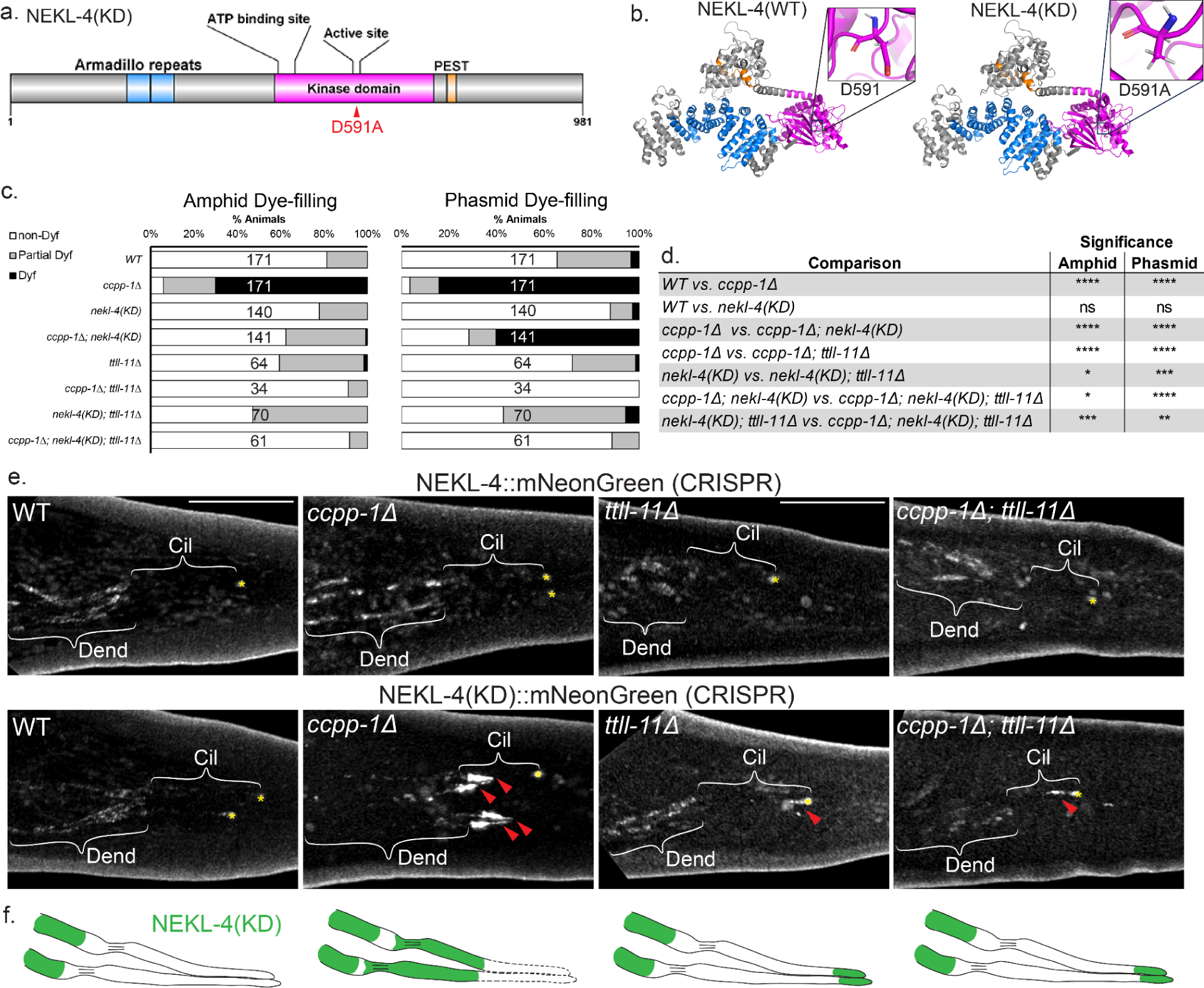
Loss of NEKL-4 kinase activity suppresses *ccpp-1Δ* dye filling defects (Dyf), and NEKL-4(KD) localizes to cilia in *ccpp-1Δ, ttll-11Δ* and *ccpp-1Δ; ttll-11Δ* mutant animals. **a.** Schematic diagram of NEKL-4 domains showing the location of the D591A active site mutation. **b.** Full-length NEKL-4 and NEKL-4(KD) predicted protein structures based on the amino-acid sequence and homolog structure information. Magenta = kinase domain, blue = armadillo repeat domain, orange = PEST sequence. The positions of the active site residue D591 and the mutant D591A are highlighted. **c.** Dye-filling of Day 2 adult animals. **d.** Significance of relevant comparisons in panel c. * indicates p ≤ 0.05, ** indicates p ≤ 0.01, *** indicates p ≤ 0.001, **** indicates p ≤ 0.0001 by Kruskal-Wallis one-way ANOVA with post hoc Dunn’s correction for multiple comparisons. **e.** Images of endogenously-tagged NEKL-4::mNeonGreen in the phasmid neurons. Scale bar = 10 µm. Cil = cilia, Dend = dendrites, red arrowhead = NEKL-4 ciliary localization, * = phasmid pore. **f.** Schematics of NEKL-4(KD) localization in WT, *ccpp-1Δ, ttll-11Δ,* and *ccpp-1Δ; ttll-11Δ* mutants. Dashed lines in the *ccpp-1Δ* panel represent ciliary degeneration.

Additionally, Dyf suppression was maintained into later adulthood in *ccpp-1Δ; nekl-4(KD)* animals, even as the Dyf phenotype worsened for *ccpp-1Δ* mutants, suggesting that loss of *nekl-4* kinase activity protects neuronal cilia from age-related degeneration (**Fig S2**). We observed some differences in Dyf suppression between the amphid and phasmid cilia.

However, with respect to other phenotypes, amphid and phasmid cilia are similar. The amphid sensory organ houses 12 sensory neurons, which cannot easily be distinguished using live confocal imaging. The phasmid cilia are structurally comparable to the amphids (Tran et al., 2023), and are easily distinguished as they are the only ciliated neurons in the posterior area of the worm, other than PQR. Thus, for further experiments requiring live imaging, the phasmid neurons were used.

Next we examined wild-type NEKL-4 and NEKL-4(KD) localization in *ccpp-1(+)* and *ccpp-1Δ* mutant backgrounds (**Fig 1e**). NEKL-4::mNG was expressed in most or all ciliated neurons and localized to dendrites, cell bodies, and axons, but was undetectable in cilia, consistent with previous results (Power et al., 2020). A similar distribution of NEKL-4::mNG was also observed in a *ccpp-1Δ* mutant background (**Fig 1e**). In the *ccpp-1(+)* background, NEKL-4(KD) resembled the NEKL-4 localization pattern, though the fluorescence was slightly dimmer. Intriguingly, in a *ccpp-1Δ* mutant background, NEKL-4(KD) exhibited a novel phenotype-localization to cilia in addition to the other parts of the neuron (**Fig 1e-f**). In hyperglutamylated *ccpp-1Δ* mutants, NEKL-4(KD) localization appeared restricted to the proximal region of the cilium. Notably, NEKL-4(KD) ectopic ciliary localization occurred specifically in this *ccpp-1Δ* Dyf-suppressed mutant, whereas the wild-type NEKL-4 protein did not show such ectopic localization. These observations suggest that microtubule glutamylation state affects NEKL-4 localization.

To test whether loss of glutamylation affected NEKL-4 distribution, we examined NEKL-4 and NEKL-4(KD) in a *ttll-11Δ* glutamylase mutant background. In both *ttll-11Δ* and *ccpp-1Δ; ttll-11Δ* mutants, NEKL-4 localization was similar to the WT *ccpp-1(+); ttll-11(+)* background (**Fig 1e**). In contrast, NEKL-4(KD) ectopically localized to the distal region of cilia of *ttll-11Δ* single and *ccpp-1Δ; ttll-11Δ* double mutants. These results are consistent with perturbed microtubule glutamylation state affecting the ciliary localization of NEKL-4(KD). Additionally, the region of the cilium where NEKL-4(KD) ectopically located varied based on hyperglutamylation (proximal region in *ccpp-1Δ* single mutant) or hypoglutamylation (distal region *ttll-11Δ* single and *ccpp-1Δ; ttll-11Δ* double mutants).

Since NEKL-4(KD) only localized to cilia in the *ccpp-1Δ* mutant background, we hypothesized that microtubule glutamylation state affects NEKL-4 localization. To test this, we examined NEKL-4 and NEKL-4(KD) localization in a *ttll-11Δ* glutamylase mutant background. In both *ttll-11Δ* and *ccpp-1Δ; ttll-11Δ* mutants, NEKL-4 localization was similar to WT (**Fig 1e**). In contrast, NEKL-4(KD) in both mutants also localized to the distal region of cilia. This indicates that perturbed microtubule glutamylation state affects the ciliary localization of NEKL-4(KD), and that the region of the cilium in which this occurs varies based on which parts of the glutamylation/deglutamylation machinery are mutated.

Since *nekl-4(KD)* suppressed the *ccpp-1Δ* dye-filling defect, we hypothesized that *nekl-4* may genetically interact with other components of the glutamylation/deglutamylation machinery. We performed dye-filling assays on double and triple mutants of *nekl-4(KD), ccpp-1Δ,* and glutamylase mutant *ttll-11Δ,* which lacks detectable ciliary glutamylation (Power et al., 2020). As published previously, *ttll-11Δ* was not Dyf in comparison to WT and suppressed the dye-filling defect of *ccpp-1Δ* (**Fig 1c-d**). We found that the *nekl-4(KD); ttll-11Δ* double mutant Dye-filling phenotype resembled the *ttll-11Δ* single mutant and that *ccpp-1Δ; nekl-4(KD); ttll-11Δ* triple mutants were similar to *ccpp-1Δ; ttll-11Δ* double mutants. However, the *nekl-4(KD); ttll-11Δ* mutants differed from the *nekl-4(KD)* mutant and *ccpp-1Δ; nekl-4(KD); ttll-11Δ* triple mutants differed from both *ccpp-1Δ; nekl-4(KD)* and *nekl-4(KD); ttll-11Δ* double mutants. Hence, the *nekl-4(KD)* mutation mildly interacts with *ttll-11Δ* with respect to dye filling.

### The nekl-4(KD) mutation does not directly affect ciliary glutamylation state

To determine if our *nekl-4(KD)* allele affected microtubule glutamylation, we measured glutamylation in the phasmid cilia. We used GT335, a monoclonal antibody that detects branch-point glutamylation and labels the doublet microtubule region of *C. elegans* cilia (Wolff et al., 1992; Kimura et al., 2010; Wloga et al., 2017; O’Hagan et al., 2017). We previously showed that *ccpp-1* deglutamylase mutants, contrary to what was expected, do not stain with GT335 in the amphid cilia, likely due to loss of doublet microtubules and ciliary degeneration (O’Hagan et al., 2017; Power et al., 2020). We found that both *nekl-4*(*my31*) and *nekl-4Δ* stain with GT335 similarly to WT, indicating that loss of *nekl-4* function did not affect glutamylation and did not suppress the glutamylation defects in *ccpp-1Δ* amphid cilia (Power et al., 2020). Similarly, *nekl-4(KD)* did not affect glutamylation in the amphid and phasmid cilia, and did not suppress the loss of GT335 staining in *ccpp-1Δ* mutants (**Fig S3**). This result is consistent with *nekl-4(KD)* suppression of *ccpp-1Δ* Dyf by a glutamylation-independent mechanism.

### The nekl-4(KD) mutation suppresses B-tubule instability in ccpp-1Δ cilia and causes other axonemal microtubule abnormalities

NEKL-4(KD) did not impact ciliary glutamylation, yet ectopically localized to abnormally-glutamylated cilia and suppressed the *ccpp-1Δ* Dyf phenotype. Therefore, we examined the microtubule ultrastructure of *nekl-4(KD)* and *ccpp-1Δ; nekl-4(KD)* amphid channel cilia (**Fig 2a-b**). The WT amphid cilium contains nine microtubule doublets composed of complete A-tubules and incomplete B-tubules and a variable amount of microtubule singlets in the center. In adult *ccpp-1Δ* mutant amphid cilia, microtubule doublets typically have open, hook-shaped B-tubules, indicating that B-tubule stability is reduced due to hyperglutamylation (O’Hagan et al., 2011).

**Figure 2.**
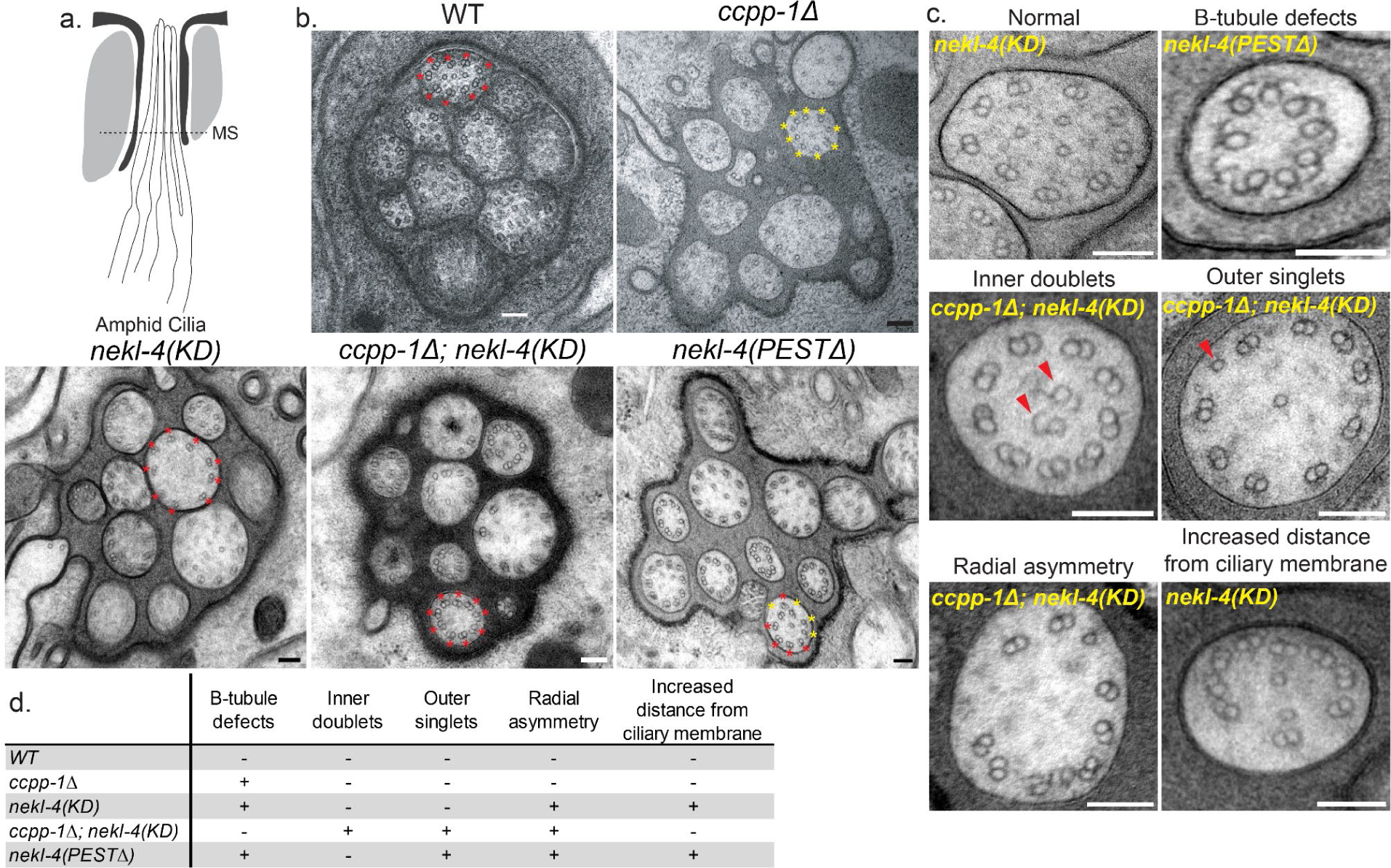
*nekl-4(KD)* mutation suppresses B-tubule instability in *ccpp-1Δ* amphid cilia. **a.** Diagram of the amphid cilia indicating the position of the cross-sections examined. MS = middle segment. **b.** Examples of microtubule doublet structure in WT, *ccpp-1Δ, nekl-4(KD), ccpp-1Δ; nekl-4(KD),* and *nekl-4(PESTΔ)* mutants. The WT and *ccpp-1Δ* panels are reproduced from O’Hagan, *et al* 2011 (O’Hagan et al., 2011) and show a male amphid. All other genotypes show a hermaphrodite amphid. Red * = normal microtubule doublet, yellow * = microtubule doublet with broken or missing B-tubule in a representative cilium. Scale bars = 100 nm. **c.** Examples of phenotypes seen in *nekl-4(KD), ccpp-1Δ; nekl-4(KD),* and *nekl-4(PESTΔ)* mutant cilia. Scale bars = 100 nm. **d.** Table of phenotypes observed in each genotype.

We rarely observed abnormal B-tubules in *nekl-4(KD)* mutants, consistent with normal localization of doublet-region exclusive kinesin-2 (**Fig S4**). In *ccpp-1Δ; nekl-4(KD)* double mutants, the number of open B-tubule doublets was reduced in comparison to *ccpp-1Δ* (**Fig 2b**). In *ccpp-1Δ; nekl-4(KD)*, we observed ectopic microtubule singlets in the doublet region and ectopic doublets in the singlet region of some cilia (**Fig 2c-d**), similar to mutants of Tubulin Code “readers’’ MAPK *mapk-15* and homodimeric kinesin-2 *osm-3* (Xie et al., 2020; Dobbelaere et al., 2023). We also observed additional ciliary defects in *nekl-4(KD), ccpp-1Δ; nekl-4(KD),* and *nekl-4(PESTΔ)* mutants, including loss of radial symmetry and increased, irregular distances between the doublets and the ciliary membrane (**Fig 2c**). In general, these defects were not as severe as those in *ccpp-1Δ* mutants and do not cause a Dye-filling phenotype.

### The nekl-4(PESTΔ) mutation causes progressive ciliary degeneration

NEKL-4 influences ciliary stability, though the mechanism by which these functions are performed remains unclear. In *Tetrahymena,* overexpression of NIMA-related kinases causes ciliary shortening (Wloga et al., 2006). In *C. elegans,* NEKL-4 overexpression (OE) causes a Dyf phenotype in a kinase activity-dependent manner, and *nekl-4* is subject to regulation via RNA editing (Power et al., 2020; Li et al., 2021a). NEKL-4 contains a PEST sequence, which is usually found in proteins whose expression levels are rapidly and tightly controlled by ubiquitin-mediated degradation (Rechsteiner and Rogers, 1996; Power et al., 2020). NEK10 also contains a PEST sequence and is regulated by ubiquitination (Porpora et al., 2018). Based on these findings, we hypothesized that NEKL-4 may be negatively regulated by the PEST motif, and that PEST deletion may result in NEKL-4 overactivity and ciliary microtubule instability. To test this hypothesis, we generated a CRISPR/Cas9 mutant with a deletion of the predicted PEST domain, which we refer to as *nekl-4(PESTΔ)* (**Fig 3a**).

**Figure 3.**
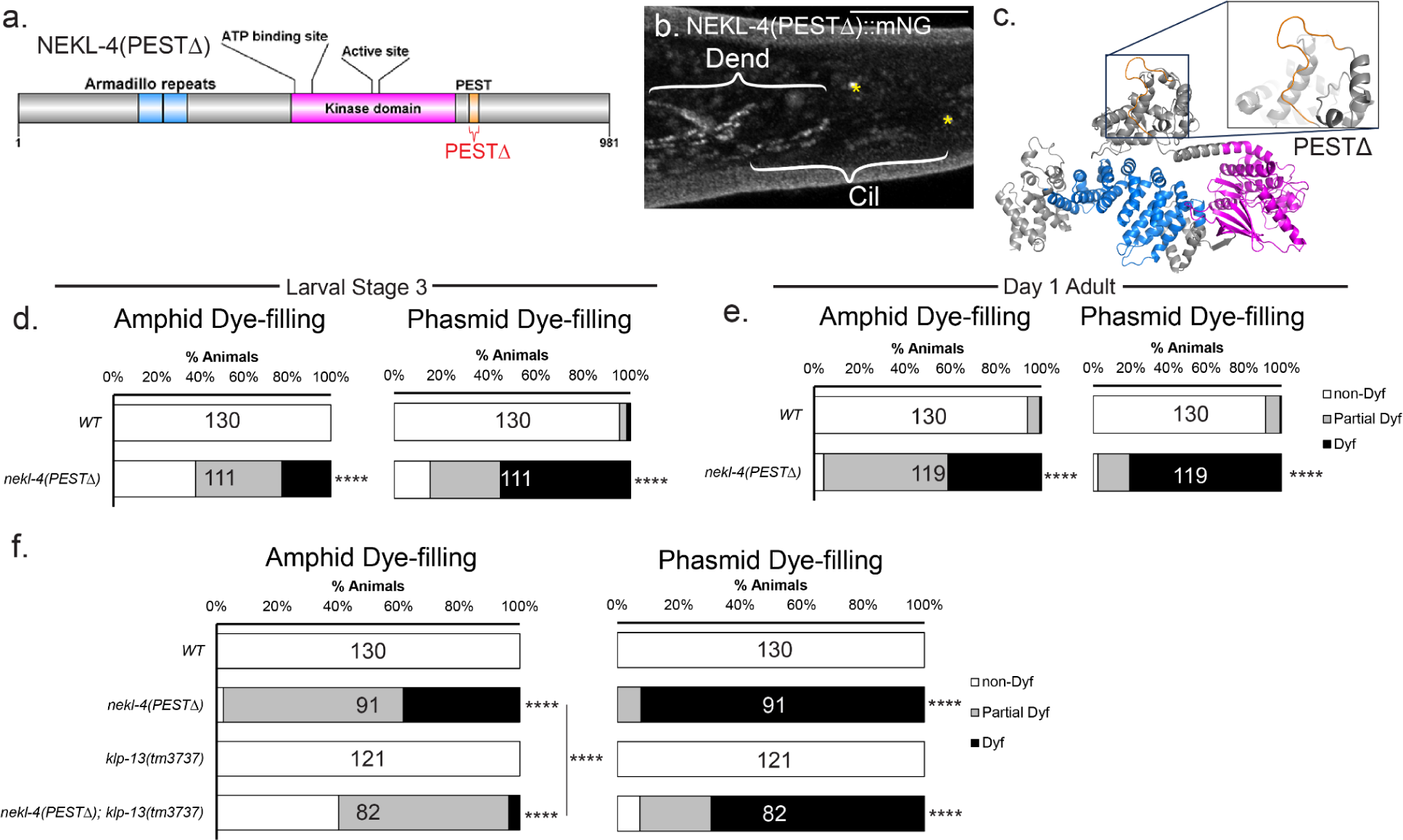
NEKL-4(PESTΔ) causes a progressive Dyf phenotype that is suppressed by mutation of kinesin-8 KLP-13/KIF19. **a.** Schematic diagram of NEKL-4 showing the location of the PESTΔ mutation. **b.** Fluorescence microscope image showing expression of NEKL-4(PESTΔ)::mNG. Scale = 10 µm. Cil = cilia, Dend = dendrites, * = phasmid pore. **c.** NEKL-4(PESTΔ) modeled protein structure. Magenta = kinase domain, blue = armadillo repeat domain, orange = residues surrounding the removed PEST sequence. The removal of the residues within the PEST domain is highlighted. **d.** Dye-filling assays of *nekl-4(PESTΔ)* L3 larvae. **** indicates p ≤ 0.0001 by Mann-Whitney test. **e.** Dye-filling assays of *nekl-4(PESTΔ)* Day 1 Adults. **** indicates p ≤ 0.0001 by Mann-Whitney test. **f.** Dye-filling assays of *nekl-4(PESTΔ); klp-13* Day 1 Adults. **** indicates p ≤ 0.0001 by Kruskal-Wallis one-way ANOVA with post hoc Dunn’s correction for multiple comparisons.

NEKL-4(PESTΔ)::mNG was still expressed by imaging the endogenous mNeonGreen tag in the strain background, and localized in a pattern similar to WT NEKL-4::mNG (**Fig 3b**). *In silico* analysis indicated that this mutation did not alter the overall structure of the protein outside of the removal of the PEST sequence (**Fig 3c**).

To determine the effect of the PEST sequence on NEKL-4 function, we performed dye-filling assays on *nekl-4(PESTΔ)* larvae stage 3 (L3) and day one adults. We hypothesized that, similar to *ccpp-1Δ, nekl-4(PESTΔ)* might cause age-dependent ciliary degeneration. Wild-type L3-stage animals have fully-developed amphid and phasmid neurons that are able to dye-fill (O’Hagan et al., 2011). At the L3 larval stage, *nekl-4(PESTΔ)* mutants were Dyf, and the severity of this defect increased as animals reached adulthood (**Fig 3d-e**). *nekl-4(PESTΔ)* phenocopied *nekl-4(OE),* consistent with dysregulated NEKL-4 causing ciliary instability. Supporting this hypothesis, we also found that *nekl-4(PESTΔ)* amphid cilia displayed more broken or defective microtubule doublets compared to WT (**Fig 2b-d**).

Our data indicated that NEKL-4 negatively regulates ciliary stability through an unknown mechanism. To explore mechanisms by which NEKL-4 may influence ciliary stability, we considered potential NEKL-4 substrates identified by phosphoproteomic analysis of NEK10-depleted cultured cells (Chivukula et al., 2020). We focused on proteins identified in cultured cells that have both a *C. elegans* ortholog and a predicted or known ciliary function in mammals or *C. elegans*: OSM-5/IFT88, DYF-5/MAK, CHE-11/IFT140, KLP-4/KIF13, and KLP-13/KIF19 (**Table S1**). OSM-5, DYF-5, and CHE-11 have been extensively characterized as regulators of *C. elegans* ciliogenesis and cilium assembly (Qin et al., 2001; Burghoorn et al., 2007; Mijalkovic et al., 2018). Kinesin-3 KLP-4 is involved in neurotransmitter trafficking; KLP-13 is a depolymerizing kinesin-8 that negatively regulates cilium length (Magaletta et al., 2019; Park et al., 2021). We therefore examined genetic interactions between *nekl-4* and these candidates. We found that the null *nekl-4Δ* allele did not modify the dye-filling phenotypes of any of the mutants, with the exception of a mild synthetic Dyf interaction with *klp-4* (**Fig S5**). *nekl-4Δ* also did not modify the dye-filling phenotypes of other genes implicated in the microtubule double region and B-tubule stability, namely *arl-13, nphp-2,* and *hdac-6* (Warburton-Pitt et al., 2014) or the dye-filling defects of ciliary kinesin-2 loss-of-function mutation *osm-3(p802)* or gain-of-function mutation *osm-3(sa125)* ((Perkins et al., 1986; Imanishi et al., 2006); **Fig S5**).

Next, we hypothesized that, since *nekl-4(PESTΔ)* mutants may be Dyf due to NEKL-4 overactivity, loss of a NEKL-4 substrate may suppress this defect. We found no genetic interaction between *nekl-4(PESTΔ)* and *osm-5, che-11,* or *dyf-5* (**Fig S6**). However, mutation of either *klp-4* or *klp-13* suppressed the Dyf defect of *nekl-4(PESTΔ)* (**Fig 3f, Fig S7**). KLP-13 is a cilia-localized microtubule-depolymerizing kinesin, and loss of *klp-13* function results in a longer ciliary axoneme (Park et al., 2021). Based on these observations, we propose that NEKL-4 negatively regulates ciliary stability via interaction with ciliary kinesins, possibly promoting KLP-13 depolymerizing activity.

### NEKL-4 is mitochondria-associated and influences mitochondrial morphology

Our results thus far indicate that NEKL-4 influences ciliary stability despite not localizing to cilia or affecting ciliary glutamylation. To begin determining the likely site of NEKL-4 function, we looked into the localization and function of mammalian CCP1 and NEK10. In neurons, loss of CCP1 results in fragmented mitochondria morphology and impaired mitochondrial transport (Gilmore-Hall et al., 2019). NEK10 is localized to mitochondria and promotes mitochondrial DNA integrity, mitochondrial respiration, and mitochondria fusion (Peres de Oliveira et al., 2020). We then sought to determine if CCPP-1 or NEKL-4 operated by similar mechanisms in *C. elegans.* To determine if NEKL-4 is mitochondria-associated, we examined localization with a mitochondrial marker. We created an extrachromosomal transgene (*osm-5p::TOMM-20::tagRFP*) that expresses an outer mitochondrial membrane marker in the ciliated neurons and examined the colocalization of TOMM-20 with NEKL-4::mNG in the phasmid dendrites (**Fig 4a**). We found that the majority of NEKL-4 protein was associated with mitochondria (**Fig 4b**), though there was a small amount of NEKL-4 that was unassociated with mitochondria. NEKL-4::mNG was co-transported with mitochondria in dendrites (**Video 1**). Mutation of the NEKL-4 active site (*nekl-4(KD)*) or *ccpp-1Δ* did not affect mitochondrial association (**Fig 4b**). To confirm the specificity of mitochondrial association, we imaged both markers in a <u>d</u>ynamin-related <u>p</u>rotein *drp-1Δ* mutant background, which is defective in mitochondrial fission and in which mitochondria are hyperfused (Labrousse et al., 1999). In the *drp-1Δ* mutant, NEKL-4 and NEKL-4(KD) remained associated with mitochondria despite the drastic change in morphology, confirming the specificity of the association (**Fig 4b**). We predict that NEKL-4 is associated with the outer mitochondrial membrane, but is not embedded within it, based on structured illumination microscopy data (**Fig 4c**) and the lack of a predicted mitochondria localization signal or transmembrane domain within the protein. NEKL-4 mitochondrial localization in *C. elegans* ciliated sensory neurons is consistent with mammalian NEK10, which is mitochondria-associated in cultured mammalian cells (Peres de Oliveira et al., 2020).

**Figure 4.**
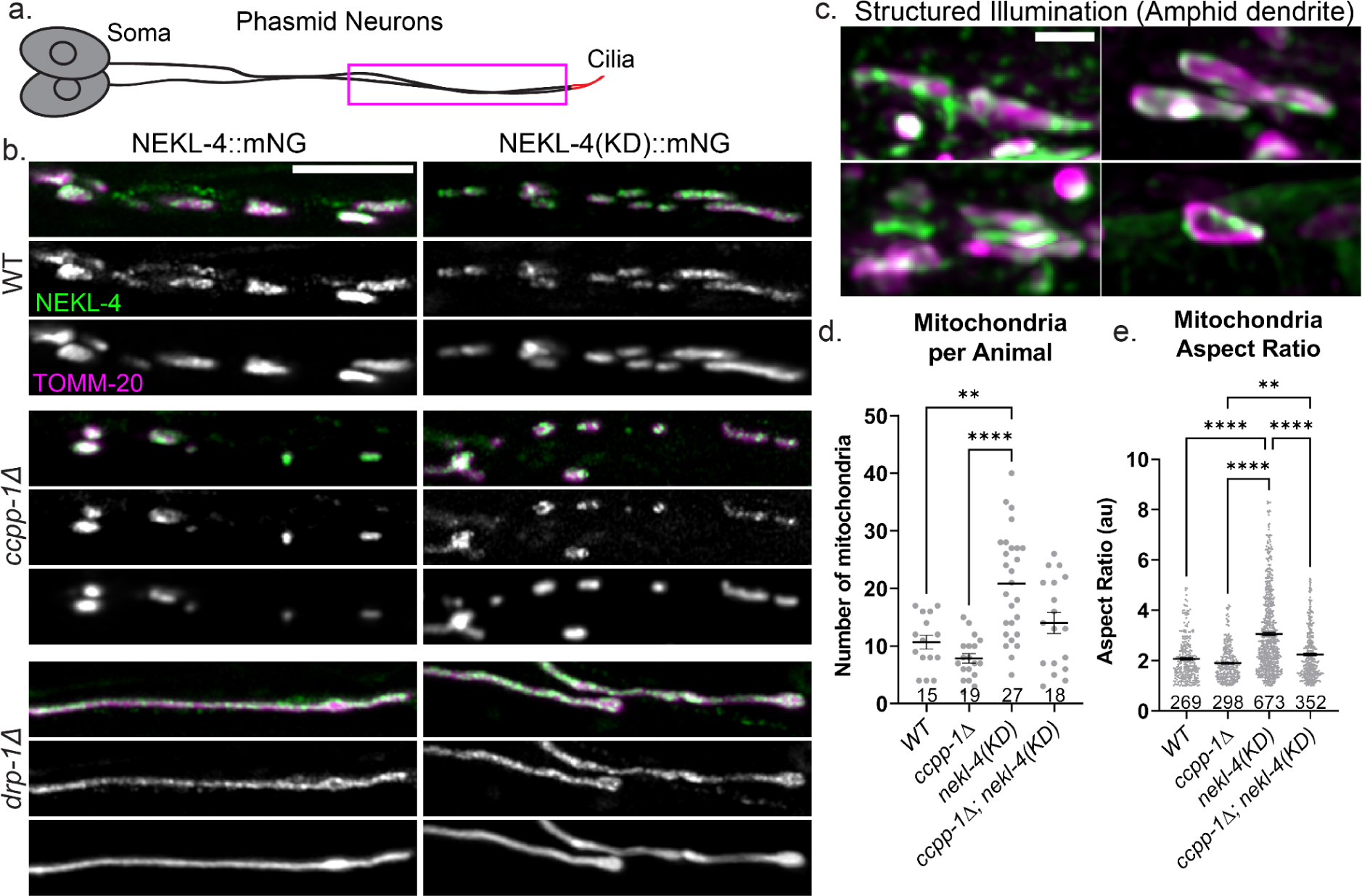
NEKL-4 is mitochondria-associated and influences mitochondrial morphology. **a.** Cartoon of one set of *C. elegans* phasmid neurons. Imaged area is boxed in magenta. **b.** Confocal images of mutants showing NEKL-4::mNG and TOMM-20::tagRFP labeling the outer mitochondrial membrane. Scale bar = 5 µm. **c.** Representative SIM images of NEKL-4::mNG and TOMM-20 colocalization on individual mitochondria in the amphid dendrites. Scale bar = 1 µm. **d-e.** Quantification of the number of mitochondria in the phasmid dendrites per animal and the aspect ratio of each mitochondrion (length/width). Mean ± SEM; ** indicates p ≤ 0.01, **** indicates p ≤ 0.0001 by Kruskal-Wallis one-way ANOVA with post hoc Dunn’s correction for multiple comparisons.

To examine mitochondrial morphological differences in *nekl-4(KD)* and *ccpp-1Δ* animals, we quantified the number and shape of mitochondria in the phasmid dendrites. We found that *ccpp-1Δ* mutant mitochondria appeared fewer in number than WT, and that *nekl-4(KD)* mutant mitochondria were more numerous (**Fig 4d**). The number of mitochondria in the *ccpp-1Δ; nekl-4(KD)* double mutant was similar to WT. Additionally, we analyzed mitochondria morphology and measured their individual aspect ratios, which is the measurement of length divided by width. *ccpp-1Δ* mutant mitochondria appeared slightly more rounded than WT, although there was no significant difference. *nekl-4(KD)* mutant mitochondria were significantly longer than WT, *ccpp-1Δ*, and *ccpp-1Δ; nekl-4(KD)* mutants (**Fig 4e**). *ccpp-1Δ; nekl-4(KD)* double mutant mitochondria were similar to WT. Based on these findings, we conclude that *nekl-4* and *ccpp-1* have opposing effects on mitochondrial morphology and number.

### NEKL-4 reduces oxidative stress in the ciliated neurons

Once we established that *nekl-4* and *ccpp-1* mutations affect mitochondria morphology and localization, we then sought to determine if CCPP-1 and/or NEKL-4 were required for mitochondria function. Mitochondria are subject to oxidative stress, which affects ciliogenesis through multiple signaling pathways (Han et al., 2021; Bae et al., 2023; Ignatenko et al., 2023). Additionally, *C. elegans* ciliated chemosensory neurons may influence longevity through oxidative stress-induced signaling, and cilia mutants are resistant to mitochondrial stress and have an increased lifespan (Apfeld and Kenyon, 1999; Fujii et al., 2004). To determine if *nekl-4(KD)* and/or *ccpp-1Δ* mutants were sensitive to oxidative stress, we performed an adult chronic paraquat assay. Paraquat (PQ) is a compound that induces oxidative stress through generation of reactive oxygen species (ROS). We plated Day 1 adult hermaphrodites on plates containing 4 mM PQ and measured the lifespan of individuals. On control plates, the lifespans of *ccpp-1Δ* and *ccpp-1Δ; nekl-4(KD)* mutants were slightly longer and shorter than WT, respectively (**Fig 5a**). On PQ, the lifespan of all mutants was extended, consistent with previous studies showing increase in longevity when exposed to mitochondrial stress (Maglioni et al., 2014; Schaar et al., 2015). On PQ, survival curves for *nekl-4(KD)* and *ccpp-1Δ; nekl-4(KD)* mutants were comparable to WT, while the lifespan of *ccpp-1Δ* mutants was significantly extended (**Fig 5a**). To further assess neuronal mitochondria stress in *nekl-4Δ* and *ccpp-1Δ* mutants, we visualized the oxidation state of mitochondria in the phasmid neurons using roGFP, a redox-sensitive GFP biosensor (Hanson et al., 2004). For this assay, we used the *nekl-4Δ* null allele to avoid interference from the mNeonGreen-tagged *nekl-4(KD)*. We found that *ccpp-1Δ, nekl-4Δ,* and *ccpp-1Δ; nekl-4Δ* mutant mitochondria all had an elevated oxidation state compared to WT, indicating that under baseline conditions, mitochondria in *nekl-4Δ, ccpp-1Δ,* and double mutants were under elevated oxidative stress (**Fig 5b**). Interestingly, though the lifespan extension phenotype of *ccpp-1Δ* under oxidative stress was suppressed by *nekl-4Δ* mutation, the same interaction was not observed when examining baseline oxidative stress. In fact, oxidation state was higher in *nekl-4Δ* mutants than *ccpp-1Δ* mutants, and double mutants were similar to *nekl-4Δ* single mutants (**Fig 5c**). Of note, total levels of roGFP were slightly elevated in *ccpp-1Δ* mutants and significantly decreased in *nekl-4Δ* and *ccpp-1Δ; nekl-4Δ* mutants, though the reason for this phenomenon is unclear (**Fig S8**). Additionally, we did not observe dendritic mitochondria in *ccpp-1Δ* or *ccpp-1Δ; nekl-4Δ* mutants, which may differ from our results in Fig 4 due to different expression levels between markers (**Fig S9**).

**Figure 5.**
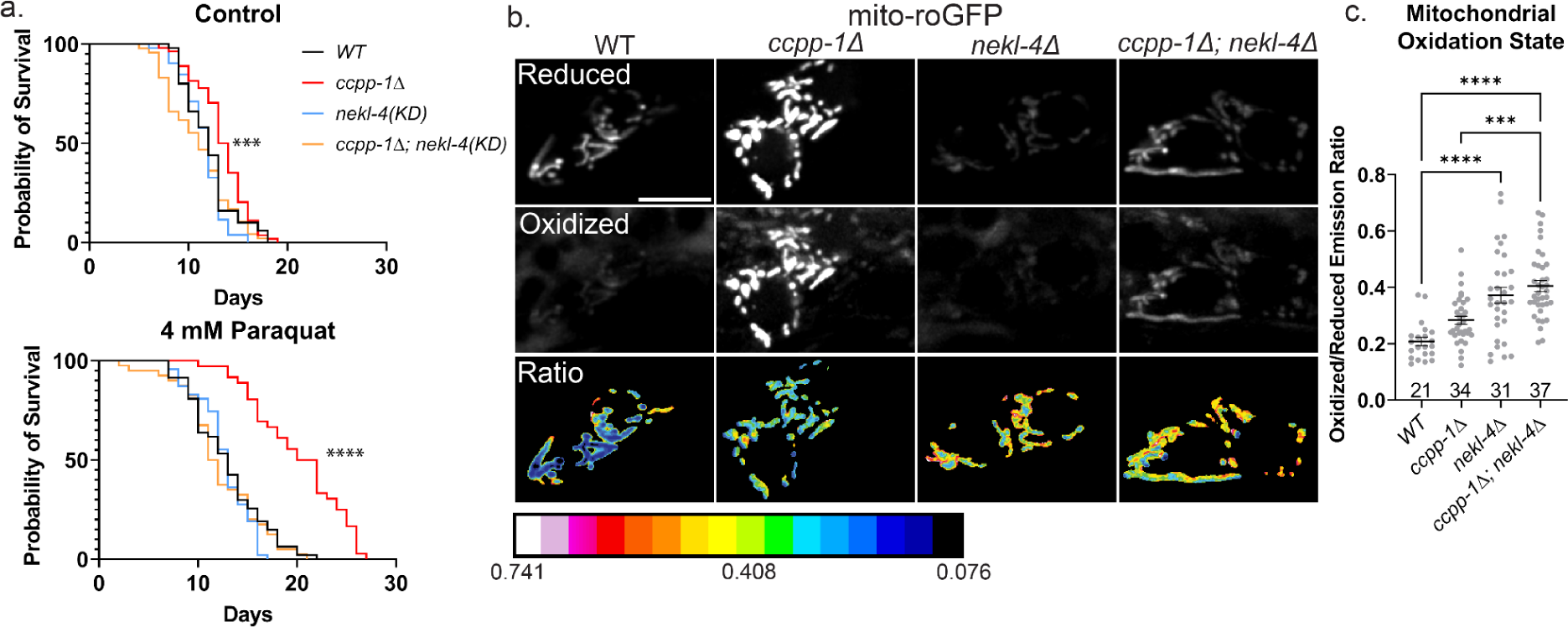
*ccpp-1Δ* and *nekl-4* mutations affect both sensitivity to oxidative stress and baseline oxidative stress in *C. elegans* ciliated sensory neurons. **a.** Survival curves of control (top) and paraquat-exposed (bottom) worms, where Day 0 is when Day 1 adult hermaphrodites were placed on plates. Control *n* = 50,54,52,47; PQ *n* = 47,36,47,40; *** indicates p ≤ 0.001, **** indicates p ≤ 0.0001 by Log-rank Mantel-Cox test. **b.** Uniformly-adjusted images of phasmid soma in the ‘reduced’ channel (top) and ‘oxidized’ channel (middle), as well as the ratio of oxidized/reduced (bottom). Scale bar = 5 µm. **c.** Quantification of oxidized/reduced emission ratios, where one point represents one set of phasmid soma. Mean ± SEM; *** indicates p ≤ 0.001, **** indicates p ≤ 0.0001 by Kruskal-Wallis one-way ANOVA with post hoc Dunn’s correction for multiple comparisons.

Overall, our results present a fascinating finding: NEKL-4 does not localize to cilia, yet has a cilia-destabilizing function as well as a mitochondrial function. Additionally, CCPP-1 is a regulator of mitochondria morphology and transport, in addition to its role in ciliary microtubule stability. NEKL-4 and CCPP-1 appear to mainly perform opposing functions in both cilia and mitochondria, and the question of how these two phenomena are related poses an exciting new direction for future studies.

## Discussion

Our previous genetic analyses indicate that NEKL-4 regulates ciliary stability in a manner downstream or independent of the glutamylation/deglutamylation machinery. In this study, we explored the NEKL-4 mechanism of action, and discovered an unexpected role in mitochondrial homeostasis in the ciliated neurons. Recent studies have revealed extensive connections between cilia stability, neurodegeneration, and oxidative stress, and our study provides evidence of NEKL-4 involvement in the intersection of these processes. We propose that NEKL-4 regulates multiple processes within the ciliated neurons via phosphorylation of multiple proteins that play diverse roles in ciliary stability and mitochondrial function.

### NEKL-4 and the Tubulin Code

NEK kinases are regulators of microtubule dynamics, particularly during the cell cycle and ciliogenesis (Li et al., 2021b). One mechanism by which NEK kinases may act is likely to involve the Tubulin Code. Tubulin post-translational modifications such as polyglutamylation influence microtubule stability and intracellular transport (O’Hagan et al., 2022; Zocchi et al., 2023). TTLL glutamylases and CCP deglutamylases act as “writers’’ and “erasers’’, while motors and microtubule-associated proteins act as “readers’’ of the Tubulin Code (Verhey and Gaertig, 2007). NEKs may regulate writers, erasers, and/or readers of the Tubulin Code. For example, NEK5 negatively regulates TTLL4 through phosphorylation, and possibly other TTLLs (Melo-Hanchuk and Kobarg, 2021). In the trypanosomatid *Crithidia fasciculata*, the NIMA-related kinase CfNek performs polyglutamylase activity, directly modifying the Tubulin Code (Westermann and Weber, 2002). Nek kinases may phosphorylate tubulin. For example, NEK1 interacts with and phosphorylates α-tubulin, with loss of function variants being a major genetic cause of amyotrophic lateral sclerosis (Mann et al., 2023). Nek kinases may also regulate microtubule-associated proteins and motors, leading to changes in activity of stability. *nekl-4* mutations do not affect glutamylation and do not modify the GT335 staining defect of *ccpp-1Δ* ((Power et al., 2020), Fig 5). While NEKL-4 does not directly modify the glutamylation state of ciliary microtubules, we propose that NEKL-4 regulates other components of the Tubulin Code. *NEKL-4(KD) ectopic ciliary localization in glutamylation mutants and NEKL-4(ΔPEST) genetic interaction with KLP-13*

Wild-type NEKL-4::mNG and NEKL-4(KD)::mNG are notably absent from cilia, raising the question of how this NEK kinase regulates ciliary stability. One clue comes from the localization of NEKL-4(KD) in glutamylation-defective mutants. In *ccpp-1Δ* and *ttll-11Δ* single mutants and *ccpp-1Δ; ttll-11Δ* double mutants, NEKL-4(KD) ectopically localizes to cilia. Though protein modeling did not reveal a significant difference in protein folding or surface electrostatic potential between NEKL-4(WT) and NEKL-4(KD) (**Fig S1**), there is a predicted reduction in cleft volume and depth in NEKL-4(KD). We hypothesize that this change in the protein-binding cleft may influence NEKL-4 interactions with binding partners as well as substrates. Kinase-dead proteins are predicted to be unable to release their substrates (He et al., 2009; Hewezi et al., 2015). If this is the case for NEKL-4(KD), then NEKL-4(KD) may be carried into cilia in *ccpp-1Δ* and *ttll-11Δ* mutants via a “stuck” substrate, such as kinesin-8 KIF19/KLP-13. KIF19 is a predicted NEK10 substrate (Chivukula et al., 2020). In *C. elegans* phasmid neurons, KLP-13 localizes to the tips of cilia, where it likely performs its microtubule depolymerizing function (Park et al., 2021). In mammalian cells, KIF19A localizes to ciliary tips and performs a dual function as both a microtubule plus-end-directed motor and a depolymerizing kinesin to regulate ciliary length (Niwa et al., 2012). Since *klp-13* mutation suppresses the ciliary degeneration of *nekl-4(PESTΔ)* mutants, we hypothesize that NEKL-4 positively regulates KLP-13 depolymerase activity, in order to negatively regulate microtubule stability (see Model in **Fig 6**).

**Figure 6.**
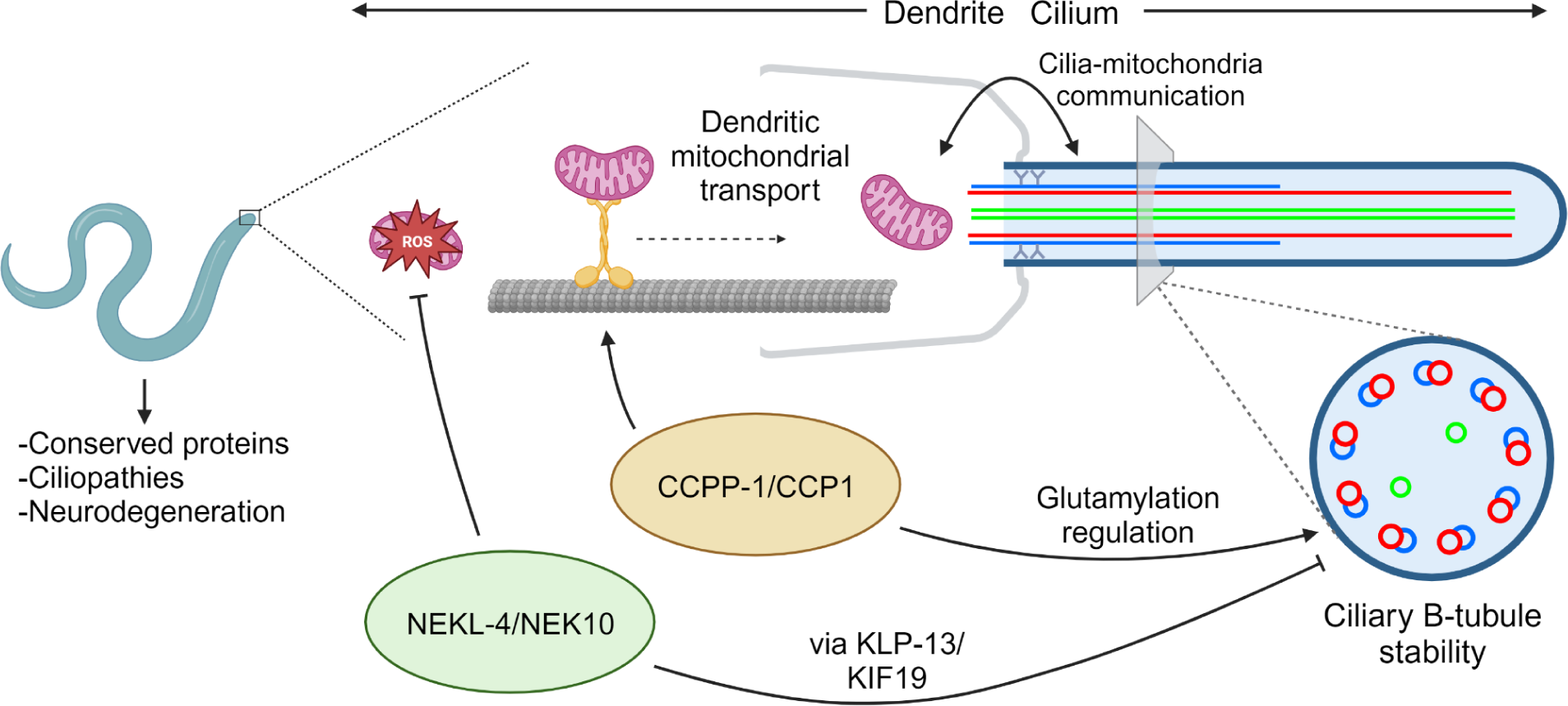
Model for NEKL-4 function in relation to cilia, mitochondria, and CCPP-1. NEKL-4 functions in the ciliated neurons to reduce mitochondrial oxidative stress and destabilize ciliary B-tubules through the action of ciliary kinesins such as KLP-13. This ciliary function opposes the B-tubule stabilizing action of CCPP-1 through a pathway distinct from glutamylation regulation. CCPP-1 also promotes dendritic transport in the ciliated neurons, specifically transport of mitochondria. Created with BioRender.com.

### A role of NEKL-4 in mediating cilia-mitochondria communication

NEKL-4(WT) did not detectably localize to cilia in any genetic backgrounds examined. Instead, NEKL-4 was mitochondria-associated and colocalized and co-transported with the TOMM-20 outer mitochondrial membrane marker. In ciliated sensory neurons, NEKL-4 modulated both oxidative stress response and mitochondrial morphology. Similar to our findings for *ccpp-1Δ*, CCP1 knockdown in cultured mammalian neurons results in mitochondria fragmentation and axonal transport defects (Magiera et al., 2018; Gilmore-Hall et al., 2019; Bodakuntla et al., 2020). Since we found a reduction in dendritic mitochondria in *ccpp-1Δ* mutants, we postulate that CCPP-1 plays a similar role in promoting mitochondrial transport in *C. elegans* ciliated neuron dendrites.

In different mammalian cell types, mitochondrial stress may promote either ciliogenesis or ciliary degeneration, but in general, increased ciliogenesis improves survival of cells when exposed to ROS (Han et al., 2021; Moruzzi et al., 2022; Ignatenko et al., 2023; Bear and Caspary, 2023; Bae et al., 2023). Cilia stability is influenced in opposing directions by mitochondrial dynamics: mitochondrial fragmentation, which increases ROS, increases ciliogenesis. Conversely, mitochondrial fusion decreases cilia length (Bae et al., 2019). In HeLa cells, NEK10 is mitochondria-localized and interacts with several mitochondrial proteins, the number of which increases in response to DNA damage-induced stress (Peres de Oliveira et al., 2020). The absence of NEK10 negatively impacts mitochondrial respiration and increases ROS, consistent with our findings that *nekl-4* mutants experience higher baseline oxidative stress in the ciliated neurons. NEK10 loss-of-function mutation results in shorter mitochondria, in contrast to *C. elegans nekl-4Δ* causing longer mitochondria, suggesting differences between NEKL-4 and its ortholog (Peres de Oliveira et al., 2020). Regardless, NEKL-4 and NEK10 appear to play evolutionarily conserved roles in inter-organelle communication between cilia and mitochondria (**Fig 6**).

Cilia communicate with other organelles to regulate diverse cellular processes. For example, ciliary-nuclear crosstalk is essential to organism development via signaling by Hedgehog (Gigante and Caspary, 2020) and other transcription factors (Nishimura et al., 2019). Cilia and the autophagy machinery share bi-directional communication to regulate ciliogenesis and proteolysis (Wiegering et al., 2019). In the nervous system, cilia are essential for modulating the creation of new synapses (Kumamoto et al., 2012), and, in the brain, cilia physically associate with axons to form synapses (Sheu et al., 2022). Cilia-mitochondria crosstalk also regulates cellular responses to mitochondrial stress (Silva and Cavadas, 2023), indicating that cilia are vital for signaling and sensing oxidative stress. NEKL-4 and CCPP-1 are expressed in the ciliated neurons (O’Hagan et al., 2011; Power et al., 2020) and appear to play a role in whole-organism oxidative stress response, leading us to hypothesize that these proteins act to mediate the communication between cilia and mitochondria, which then influences the signals sent to the rest of the animal by the ciliated neurons in response.

Our work offers insights into the multi-functional protein NEKL-4 and its roles in cilia and mitochondria. NEKL-4 both modulates ciliary stability and promotes mitochondrial homeostasis, similar to mammalian NEK10. Our work adds to the growing number of proteins that directly or indirectly regulate the Tubulin Code. Additionally, proteins that have ciliary functions may also influence processes in other subcellular locations, such as mitochondria. Further exploration of NEKL-4/NEK10 and other Tubulin Code-related proteins will provide insight into the communication between cilia and other organelles as well as the effects this communication may have on neurodegeneration and ciliopathies.

**Video 1. NEKL-4 and mitochondria are co-transported in dendrites.** Retrograde movement is shown beginning at the amphid distal dendrite. Scale bar = 10 µm. Played back at 10fps.

## Materials and Methods

### *C. elegans* strains and maintenance

Nematodes were cultured on Nematode Growth Media (NGM) agar plates containing a lawn of OP50 *E. coli* and incubated at 20°C, as previously described (Brenner, 1974). Strains used are listed in **Table S2**. Strains used in supplemental material are listed in **Table S3**. Genetic sequence information was obtained from WormBase (Davis et al., 2022).

### CRISPR constructs

For creation of the *nekl-4(KD)* and *nekl-4(PESTΔ)* CRISPR mutants (*my66* and *my120*), we followed the Mello Lab protocol (Dokshin et al., 2018) and used ssOligos as donors. The guide sequences were designed using CRISPOR (Haeussler et al., 2016) and silent mutation sites for PAM modification and diagnosis were located using WatCut silent mutation scanning. For *nekl-4(KD),* we modified the predicted active site (D591) to alanine, which is expected to eliminate NEKL-4 kinase activity without altering the gross structure of the protein. For *nekl-4(PESTΔ),* we deleted the predicted PEST proteolytic degradation domain (17AA; 748–764, KIDESPSSLNSSTSSYK) using an ssOligo with the sequence encoding those residues removed. This mutant is recessive for the Dyf phenotype. Both mutants were generated in the *nekl-4(my51[nekl-4::mNG])* background. CRISPR-generated strains were outcrossed at least 3X before use. For genotyping genetic crosses into strains not in the *nekl-4(my51[nekl-4::mNG])* background, primers for NEKL-4::mNG were used for diagnosis of *nekl-4(KD)* and *nekl-4(PESTΔ)*.

We also followed the same protocol to recreate the *pha-1(e2123)* temperature-sensitive point mutation in a *nekl-4Δ* mutant background, since the genes are the same linkage group. We validated that our version of the allele, *pha-1(my82[pha-1(e2123)+SnaBI]),* is temperature-sensitive before constructing strains or performing experiments with the mutant. Additionally, we inserted a silent mutation to create an additional SnaBI restriction enzyme cut site for diagnosis and differentiation between our *my82* mutation and the original *e2123* allele.

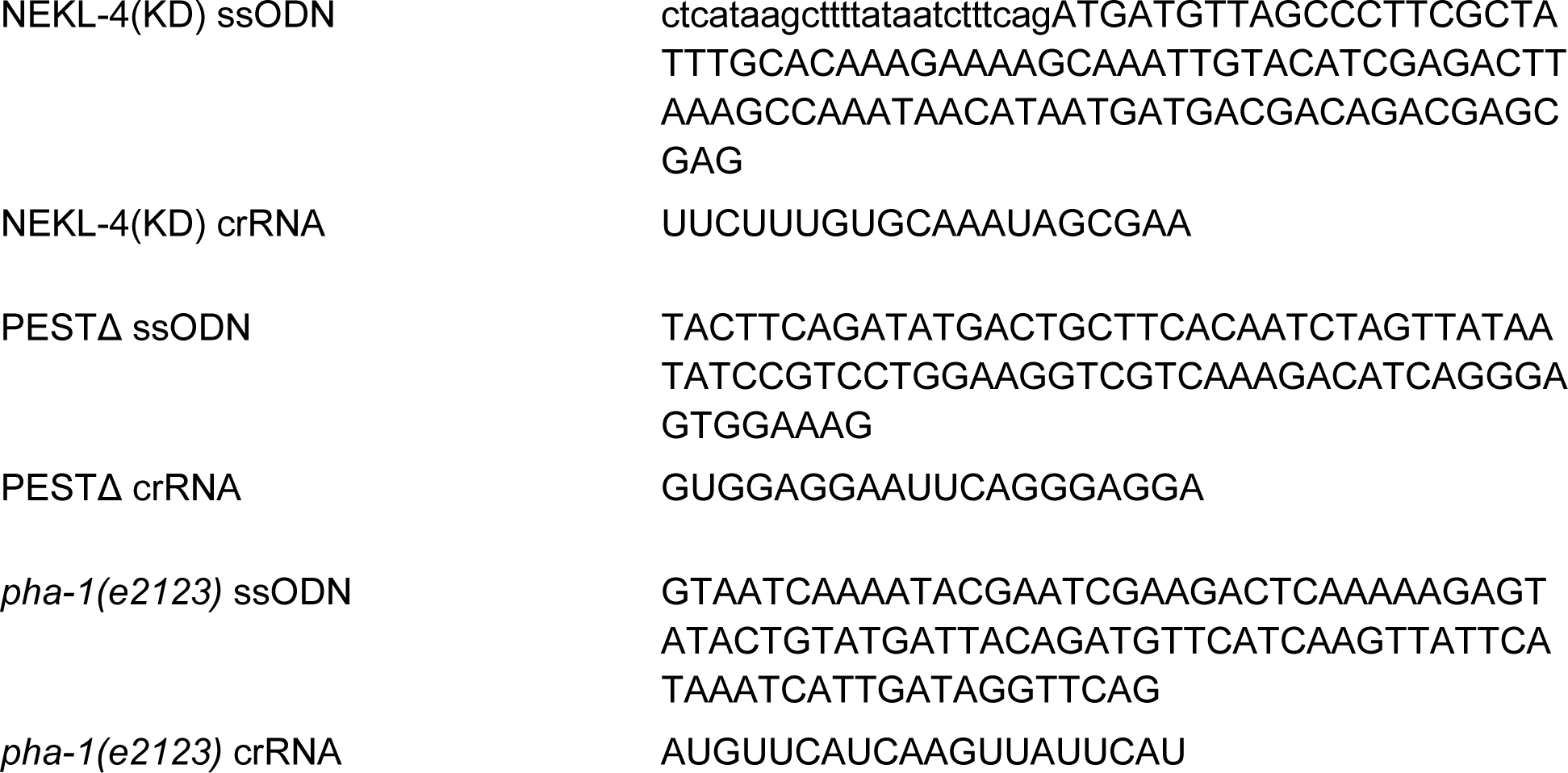

### Plasmid construction

For creation of pKP06 (*osm-5p::tomm-20::tagRFP*), the *tomm-20* coding sequence was fused to a 241bp promoter from *osm-5,* which drives expression in all ciliated neurons. Both fragments were amplified from genomic DNA using primers with homology to pPD95.75 containing *tagRFP* or the other fragment. The stop codon for *tomm-20* was removed using the reverse primer. pPD95.75 was amplified using primers with homology to *osm-5p* or *tomm-20.* Fragments were joined using Gibson assembly.

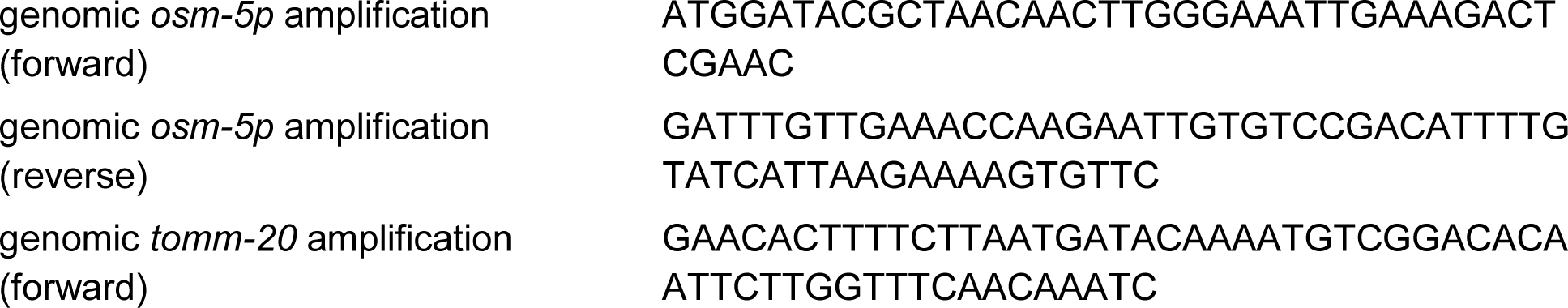
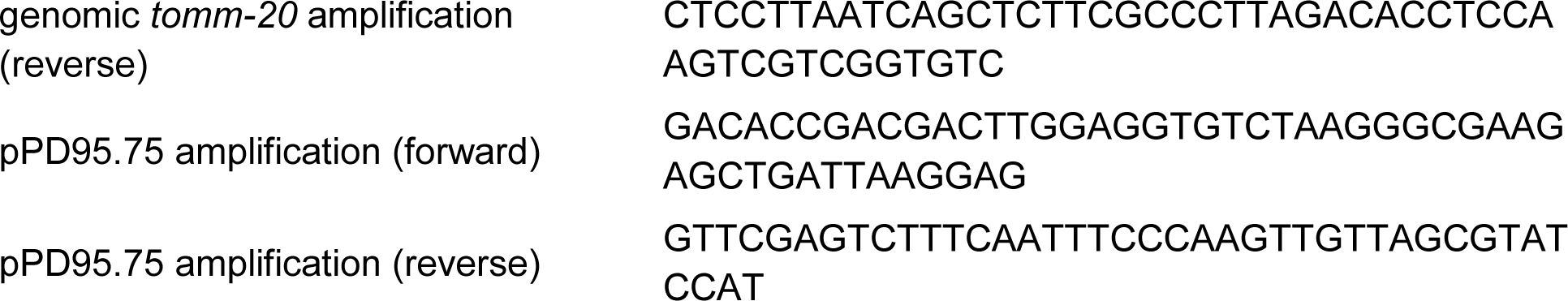

For creation of pKP07 (*osm-5p::mito-roGFP*), the previously-described *osm-5* promoter was amplified from genomic DNA using primers with homology to pOR823 containing *dat-1p::mito-roGFP.* pOR823 was amplified using primers with homology to *osm-5p.*The *dat-1* promoter was not included in amplification of the backbone. Fragments were joined using Gibson assembly.

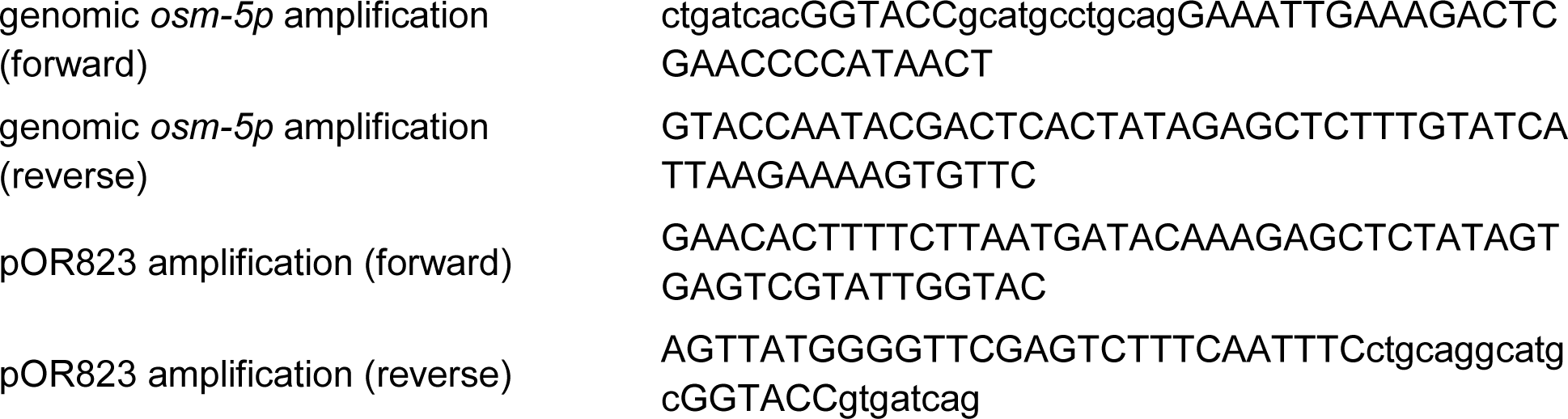

### Dye-filling assays

We performed dye-filling assays using the protocol described in (Power et al., 2020). Prior to performing dye-filling assays, healthy, non-crowded plates containing many gravid hermaphrodite animals were washed and bleached to synchronize the worms. Worms at desired ages were washed from the plate with 1mL M9 buffer and transferred to 1.5mL tubes. A stock solution of 2.5mg/mL DiI (Thermo Fisher Cat. D282) was added to a 1:250 dilution and worms were incubated with gentle shaking for 50 min at room temperature. Prior to scoring, worms were briefly spun at 2000 RPM and plated on new seeded NGM plates for approximately 1 hr to excrete excess dye. The presence and intensity of dye in the amphid and phasmid neurons was visually scored using a dissecting microscope. Worms scored as non-Dyf were similar in brightness to wild type, and Dyf worms had no staining. Partial Dyf worms had dye present, but distinguishably less bright than wild type. Kruskall-Wallis one-way ANOVA analysis (where non-Dyf=2, partial Dyf=1. and Dyf=0) and posthoc Dunn’s multiple comparison test or Mann-Whitney test were performed in Prism (Graphpad Software).

### Tertiary structure prediction and model evaluation

We predicted the structure of NEKL-4 WT and its mutants using the I-TASSER server (Yang and Zhang, 2015). In I-TASSER structural templates were first identified from the PDB database using multiple threading alignment approaches. The full-length structure models were then constructed by iterative fragment assembly simulations. The top five predicted models were rigorously evaluated by several model quality assessment programs including VERIFY3D (Lüthy et al., 1992), VoroMQA (Olechnovič and Venclovas, 2017), ProsaWEB (Wiederstein and Sippl, 2007), and ProQ3 (Uziela et al., 2016). These programs employ a variety of methods inclusive of deep learning approaches to evaluate the quality of the models. The final models were chosen by selecting the best evaluation profiles and their correlation with known functional features.

### Surface electrostatic profiles of NEKL-4 and Kinase Dead mutant

PyMOL Version 2.0 (Schrödinger LLC, New York, NY) was used to visualize the protein models and its APBS plugin (Holst and Saied, 1993) to obtain the surface electrostatic profiles of both NEKL-4 and the Kinase Dead mutant.

### Cleft Analysis of NEKL-4 and Kinase Dead mutant

PDBsum (Laskowski, 2007) was utilized for visualization and analysis of both NEKL-4 and the Kinase Dead mutant’s clefts.

### Antibody staining

We performed immunofluorescence microscopy using the protocol described in Power *et al* (Power et al., 2020). We synchronized animals by bleaching and fixed animals as Day 1 adults. Fixation was accomplished by washing animals from 3 NGM plates using M9 buffer, then washing animals with several changes of M9 over one hour. Worms were chilled on ice before washing in ice-cold Ruvkun buffer (80 mM KCl, 20 mM NaCl, 10 mM EGTA, 5 mM spermidine-HCl, 15 mM PIPES, pH 7.4 and 25% methanol) plus 2% formaldehyde in 1.5ml centrifuge tubes. The tubes were immersed in liquid nitrogen and melted under tap water to crack the worms’ cuticles. Worms were then washed twice with Tris-Triton buffer (100 mM Tris-HCl, pH 7.4, 1% Triton X-100 and 1 mM EDTA), suspended in Tris-Triton buffer + 1% β-mercaptoethanol, and incubated overnight at 37°C. The next day, worms were washed with 1X BO_3_ buffer (50 mM H_3_BO_3_, 25 mM NaOH) + 0.01% Triton, and suspended in 1X BO_3_ + 0.01% Triton buffer + 10 mM DTT for 15 minutes with gentle agitation at room temperature. Worms were then washed with 1X BO_3_ buffer (50 mM H_3_BO_3_, 25 mM NaOH) + 0.01% Triton, and suspended in 1X BO_3_ + 0.01% Triton buffer + 0.3% H_2_O_2_ for 15 minutes with gentle agitation at room temperature. After washing once with 1X BO_3_ + 0.01% Triton buffer, worms were washed for 15 minutes in Antibody buffer B (1X PBS, 0.1% BSA, 0.5% Triton X-100, 0.05% sodium azide, 1mM EDTA) with gentle agitation at room temperature. Fixed worms were stored in Antibody buffer A (1X PBS, 1% BSA, 0.5% Triton X-100, 0.05% sodium azide, 1mM EDTA) at 4°C for up to one month before antibody staining. Animals were stained overnight at room temperature with a 1:600 final dilution (in Antibody Buffer A) of GT335 (Adipogen Cat. AG-20B-0020-C100), a monoclonal antibody which binds the branch point of both monoglutamylated and polyglutamylated substrates (Wolff et al., 1992). Stained worms were washed with several changes of Antibody B Buffer with gentle agitation at room temperature over several hours. After rinsing with Antibody Buffer A, Alexa-fluor 568-conjugated donkey anti-mouse secondary antibody (Invitrogen Cat. A10037) was added at a final dilution of 1:2500 and incubated for 2 hours at room temperature with gentle agitation. Worms were then washed with several changes of Antibody Buffer B over several hours before mounting on 10% agarose pads for imaging.

### Confocal imaging

Live animals were anesthetized with 10 mM levamisole and mounted on 10% agarose pads for imaging at room temperature. Confocal imaging was performed with a Zeiss LSM 880 inverted microscope with an Airyscan superresolution module using a LSM T-PMT detector and ZenBlack software (Carl Zeiss Microscopy, Oberkochen, Germany). Laser intensity was adjusted to avoid saturated pixels. Images were acquired using a 63x/1.4 Oil Plan-Apochromat objective in Airyscan or Airyscan Fast mode and deconvolved using Airyscan processing. Image files were imported into Fiji/ImageJ with the BioFormats Importer plugin for linear adjustment of contrast and creation of maximum intensity projections.

### Widefield imaging

Nematodes were mounted as above. Widefield images were acquired on a Zeiss Axio Observer with Colibri 7 LEDs and ZenBlue software (Carl Zeiss Microscopy, Oberkochen, Germany) using a Photometrics Prime 95B sCMOS camera (Teledyne Photometrics, Tucson, AZ). A 63x/1.4 Oil Plan-Apochromat objective was used for imaging.

### Transmission electron microscopy

Three strains, PT3471 (*nekl-4(KD)*), PT3756 (*nekl-4(PESTΔ)*) and PT3593 (*ccpp-1Δ; nekl-4(KD)*), were grown for 4 generations on seeded OP50 agarose plates at 20°C, avoiding starvation at any time. Young adult worms were fixed by high pressure freezing/freeze substitution (HPS/FS) (Hall et al., 2012) with a fixative of 2% osmium tetroxide, 0.1% uranyl acetate, 2% H_2_O in acetone, for a week at −90°C, then slowly warming stepwise to −60°C, −30°C, and 0°C. Samples were washed 4 times in 100% cold acetone, infiltrated in 5 changes into EMbed812 resin (Electron Microscopy Sciences Cat. 70020) over 2 days and cured at 60°C.

Twelve PT3471 (*nekl-4(KD)*), six PT3756 (*nekl-4(PESTΔ)*) and 17 PT3593 (*ccpp-1Δ; nekl-4(KD)*) young adult hermaphrodites were collected either at 70 nm, 90 nm, 150 nm or 200 nm transverse serial sections on Pioloform/Formvar carbon-coated wide slot grids from an RMC Ultramicrotome PowerTome PT XL microtome (Böckeler Instruments, Tucson, AZ) starting nose tip to amphid base covering about 25-40 µm of the amphid organ. Sections were post-stained in 2% uranium acetate for 20-40 min and in 1:5 dilution aqueous Reynolds lead stain for 5 min. Most animals were imaged separately for the left and right amphids from the pore to the base of the amphid with Digital Micrograph software on a Hitachi H-7500 TEM (Hitachi, Chiyoda City, Tokyo, Japan) with ATM CD camera, covering serial thin sections. In order to obtain electron tomograms, other animals were also viewed using SerialEM software on a JEM-1400Plus TEM (Jeol Ltd., Akishima, Tokyo, Japan) with Gatan Orius SC1000B bottom mount digital camera (Gatan Inc., Pleasanton, CA), using serial thick sections. Serial amphid image stacks were aligned with TrakEM2, then processed and re-aligned with Etomo and 3dmod software (Mastronarde Group, University of Colorado, Boulder, CO) to produce tomograms.

### Structured illumination microscopy (SIM)

Live Day 1 adult animals carrying *osm-5p::tomm-20::tagRFP* and endogenous *NEKL-4::mNG* were mounted as described above and imaged using a Zeiss Elyra 7 with Lattice SIM² (Carl Zeiss Microscopy, Oberkochen, Germany). We acknowledge Jessica Shivas and Nancy Kane for assistance with SIM imaging.

### Automated mitochondrial morphology quantification

Images of animals carrying *osm-5p::tomm-20::tagRFP* were acquired using the Zeiss LSM 880 confocal microscope as described above and maximum intensity projections were used for analysis. Mitochondria Analyzer, a Fiji/ImageJ macro, was used to quantify mitochondria morphology and number in phasmid dendrites (Chaudhry et al., 2020). Images were converted to 8-bit and cell bodies were trimmed. Settings used for batch 2D thresholding were as follows: Rolling 1.25 microns; max slope 1.8; gamma 0.8; block size 1.15 microns; C-Value 5. Batch 2D analysis was then performed on a per-mito basis. After automatic analysis, data were cleaned by eliminating all mitochondria with >1 branch and/or ≠2 endpoints, since mitochondria in the dendrites are unbranched and results indicating branches are likely an artifact of using Z-projected images for analysis. Mitochondria <0.3 µm were also eliminated as they are likely background noise or artifacts not eliminated by thresholding. Outliers were removed with the ROUT method (Q = 0.1%) and Kruskall-Wallis one-way ANOVA analysis and posthoc Dunn’s multiple comparison test were performed in Prism (Graphpad Software).

### Chronic paraquat survival assay

The adult chronic paraquat survival assay was performed as in Schaar *et al* (Schaar et al., 2015). Worms were synchronized by bleaching and Day 1 adults were plated on NGM plates containing 100 µM 5-Fluoro-2′-deoxyuridine (FuDR) (Sigma Cat. F0503) with or without 4 mM methyl viologen dichloride hydrate (paraquat) (Sigma Cat. 856177). FuDR was used to inhibit the development of progeny, since paraquat causes an increased rate of internal hatching that may prematurely kill the worms. Control and paraquat plates were seeded with 150 µL of 10X concentrated OP50 and allowed to dry before use. Adult worms were transferred to new plates every seven days until there were no surviving animals. Worms were considered dead when unmoving and unresponsive to multiple touch stimuli. Worms that died during transfer or by crawling off the plate were censored. Data were visualized as Kaplan-Meier survival curves and analyzed using a log-rank Mantel-Cox test performed in Prism (Graphpad Software).

### Mitochondrial redox state analysis

Nematodes carrying a mitochondria-targeted redox sensor expressed in the ciliated neurons (*osm-5p::mito-roGFP*) were mounted as above and imaged using the Zeiss LSM 880 confocal microscope. The mitochondria targeting signal for this biosensor was taken from aspartate aminotransferase. roGFP (reduction-<u>o</u>xidation-sensitive GFP) is a modified GFP that contains cysteines at certain residues that create two excitation peaks corresponding to different oxidation states (Hanson et al., 2004). By measuring the ratio of emissions at each wavelength, the relative oxidation state of mitochondria can be monitored. Phasmid cell bodies of Day 1 adult worms were imaged with 405 nm (oxidized) and 488 nm (reduced) excitation. Only mitochondria close to the coverslip were imaged to minimize differences in emission intensity caused by differences in the depth of the cells within the worm. For analysis, maximum intensity projections were created from 3-5 slices per worm. Channels were split and the 488 nm channel was duplicated and used to create ROIs including only mitochondria by thresholding. The mean fluorescence intensity of each channel within the ROI was measured, and the 405 nm intensity was divided by the 488 nm intensity to obtain the final emission ratio. To generate the ratio images, original images were converted to 32-bit and the 405 nm channel was divided by the 488 nm channel using the ImageJ image calculator and displayed using the “16 colors” LUT. Outliers were removed with the ROUT method (Q = 0.1%) and Kruskall-Wallis one-way ANOVA analysis and posthoc Dunn’s multiple comparison test were performed in Prism (Graphpad Software).

### Supplemental Results

**Supplemental Table 1.** Candidate interactors of *nekl-4* and their functions. Adapted from (Power et al., 2020)

| Gene | Function |
| --- | --- |
| *osm-5* | IFT-B component |
| *dyf-5* | IFT cargo and component handover |
| *che-11* | IFT-A component |
| *klp-4* | Neurotransmitter trafficking kinesin |
| *klp-13* | Depolymerizing kinesin |

**Supplemental Table 2.**
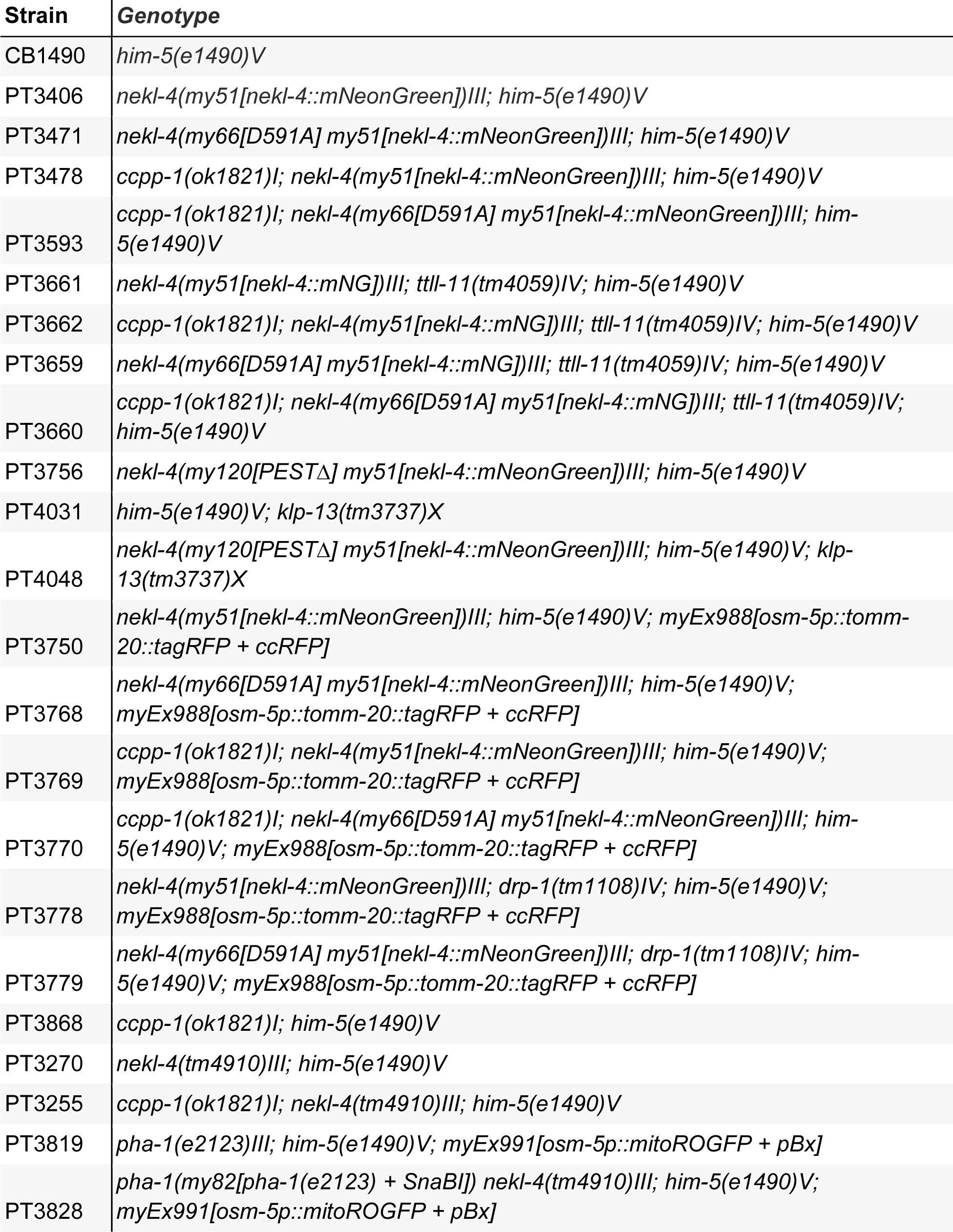
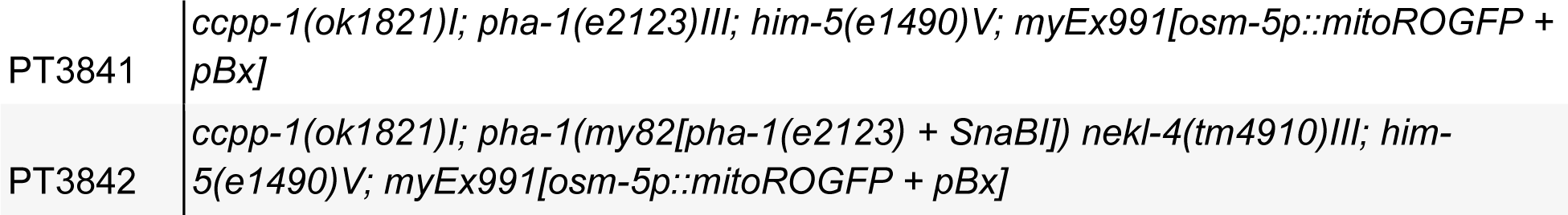
Strains used in this study.

**Supplemental Table 3.** Strains used in the supplemental material.

| Strain | Genotype |
| --- | --- |
| GOU1810 | *casIs586[kap-1p::kap-1::GFP + pRF4]* |
| PT4050 | *nekl-4(my66[D591A] my51[nekl-4::mNeonGreen])III; casIs586[kap-1p::kap-1::GFP + pRF4]* |
| CB3330 | *che-11(e1810)V* |
| PT3801 | *nekl-4(tm4910)III; che-11(e1810)V* |
| PT3298 | *dyf-5(ok1170)I; him-5(e1490)V* |
| PT3305 | *dyf-5(ok1170)I; nekl-4(tm4910) III; him-5(e1490)V* |
| PT2241 | *him-5(e1490)V; osm-5(m184)X* |
| PT3840 | *nekl-4(tm4910)III; him-5(e1490)V; osm-5(m184)X* |
| PT2815 | *arl-13(gk513)I; him-5(e1490)V* |
| PT3500 | *arl-13(gk513)I; nekl-4(tm4910)III; him-5(e1490)V* |
| PT2145 | *nphp-2(gk653) him-5(e1490)V* |
| PT3511 | *nekl-4(tm4910)III; nphp-2(gk653) him-5(e1490)V* |
| PT2816 | *hdac-6(ok3203)IV; him-5(e1490)V* |
| PT3504 | *nekl-4(tm4910)III; hdac-6(ok3203)IV; him-5(e1490)V* |
| PT1974 | *osm-3(p802)IV; him-5(e1490)V* |
| PT3739 | *osm-3(my115[sa125(G444E)])IV; him-5(e1490)V* |
| PT3347 | *nekl-4(tm4910)III; osm-3(p802)IV; him-5(e1490)V* |
| PT3771 | *nekl-4(tm4910)III; osm-3(my115[sa125(G444E)])IV; him-5(e1490)V* |
| PT3773 | *nekl-4(my66[D591A] my51[nekl-4::mNeonGreen])III; osm-3(my115[sa125(G444E)])IV; him-5(e1490)V* |
| PT974 | *him-5(e1490)V; klp-4(my5)X* |
| PT1254 | *him-5(e1490)V; klp-13(my7)X* |
| PT3591 | *nekl-4(tm4910)III; him-5(e1490)V; klp-4(my5)X* |
| PT3592 | *nekl-4(tm4910)III; him-5(e1490)V; klp-13(my7)X* |
| PT3982 | *nekl-4(my120[PESTΔ] my51[nekl-4::mNeonGreen])III; che-11(e1810)V* |
| PT4009 | *dyf-5(ok1170)I; nekl-4(my120[PESTΔ] my51[nekl-4::mNeonGreen])III; him-5(e1490)V* |
| PT4010 | *nekl-4(my120[PESTΔ] my51[nekl-4::mNeonGreen])III; him-5(e1490)V; osm-5(m184)X* |
| PT3981 | *nekl-4(my120[PESTΔ] my51[nekl-4::mNeonGreen])III; him-5(e1490)V; klp-4(my5)X* |

**Supplemental Figure 1.**
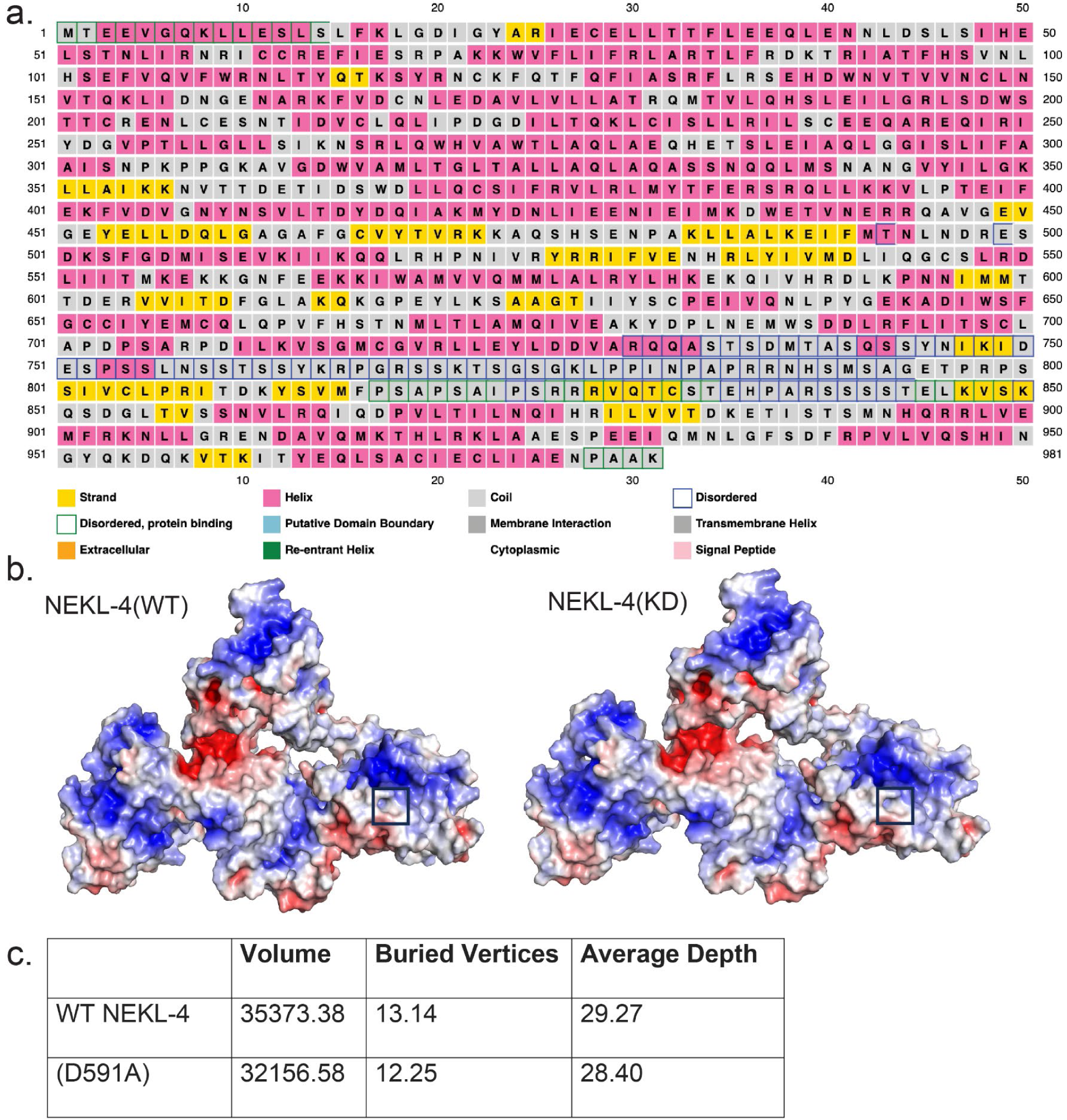
I*n silico* modeling of NEKL-4 and NEKL-4(KD) **a.** Secondary structure prediction and prediction of intrinsically disordered regions of NEKL-4. **b.** Surface Electrostatic Potential of NEKL-4 and NEKL-4(KD). Blue = Positive, Red = Negative (−5 to +5 kT/e). Boxed area is residue D591 or D591A. **c.** Cleft analysis of NEKL-4 and NEKL-4(KD).

**Supplemental Figure 2.**
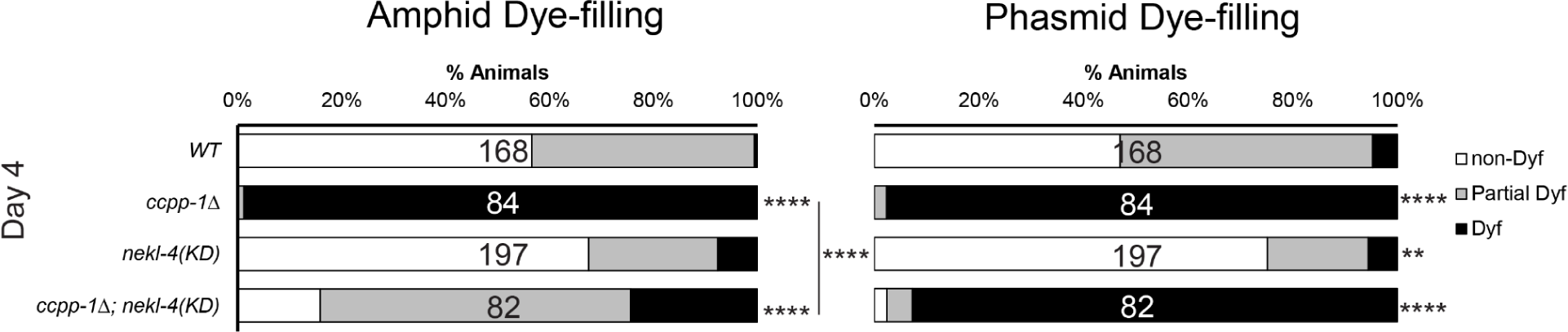
*nekl-4(KD)* mutation continues to suppress the *ccpp-1Δ* dye-filling defect into later adulthood. ** indicates p ≤ 0.01, **** indicates p ≤ 0.0001 by Kruskal-Wallis one-way ANOVA with post hoc Dunn’s correction for multiple comparisons.

**Supplemental Figure 3.**
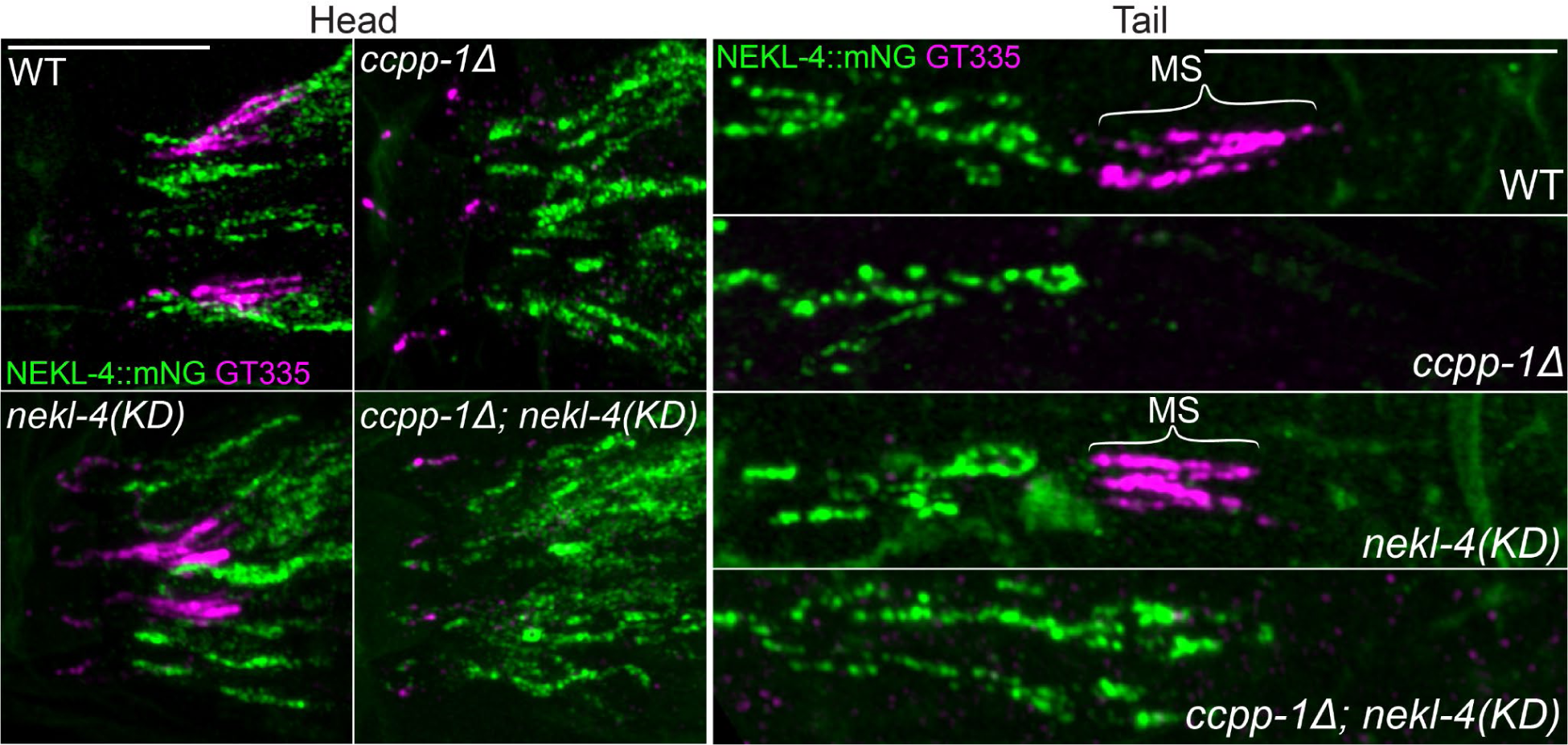
Loss of NEKL-4 kinase activity does not suppress the loss of GT335 staining in *ccpp-1Δ* mutant amphid and phasmid cilia. Images of endogenously-tagged NEKL-4::mNeonGreen and GT335, a monoclonal antibody that detects mono- and polyglutamylation, in the amphid, labial, and cephalic cilia. Scale bars = 10 µm. MS = middle segment.

**Supplemental Figure 4.**
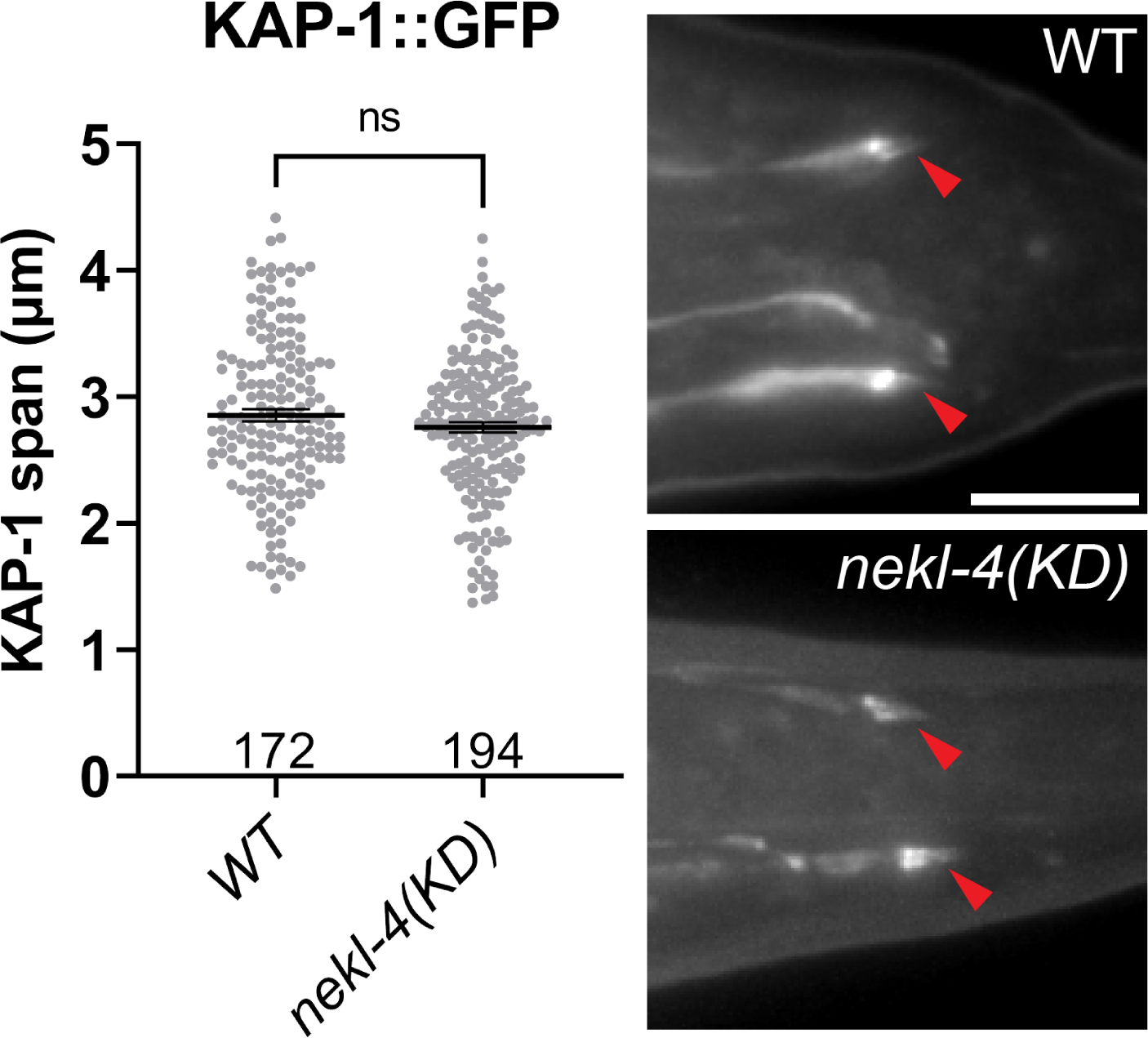
KAP-1 span is not altered in *nekl-4(KD)* mutants. Left: quantification of KAP-1 span in the phasmid cilia. Not significant by Mann-Whitney test. Right: Examples of KAP-1::GFP localization. Red arrowhead = phasmid cilia. Scale bar = 10 µm.

**Supplemental Figure 5.**
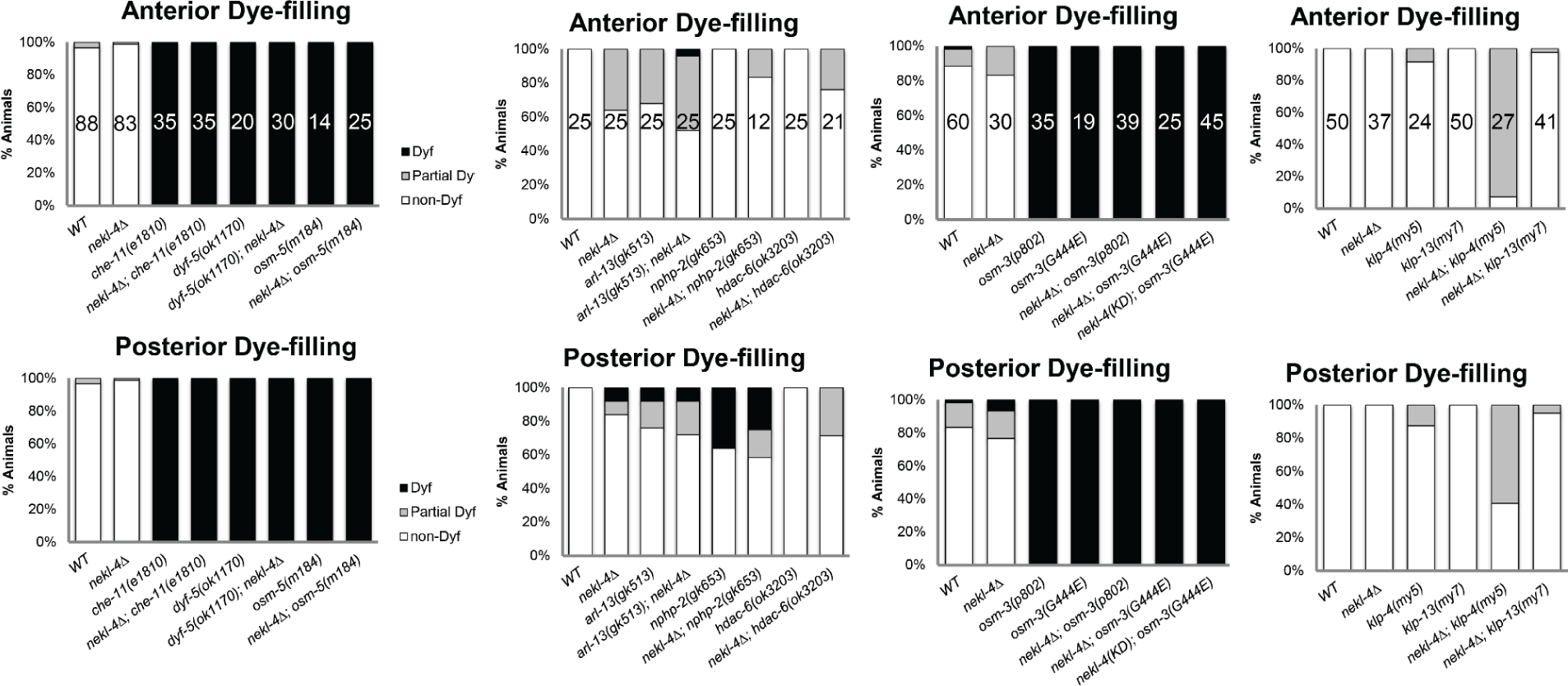
Negative dye-filling interactions for *nekl-4Δ*.

**Supplemental Figure 6.**
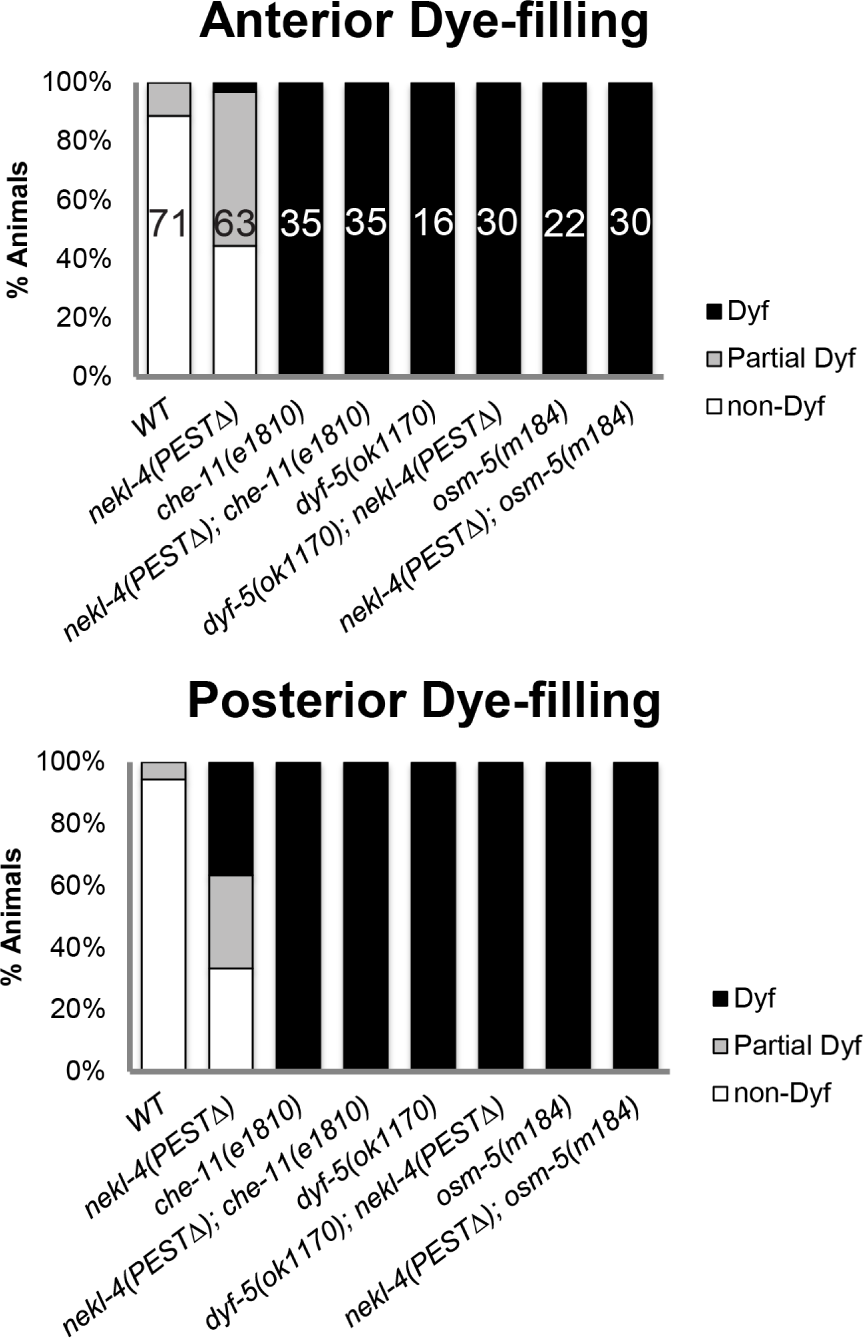
Negative dye-filling interactions for *nekl-4(PESTΔ)*

**Supplemental Figure 7.**
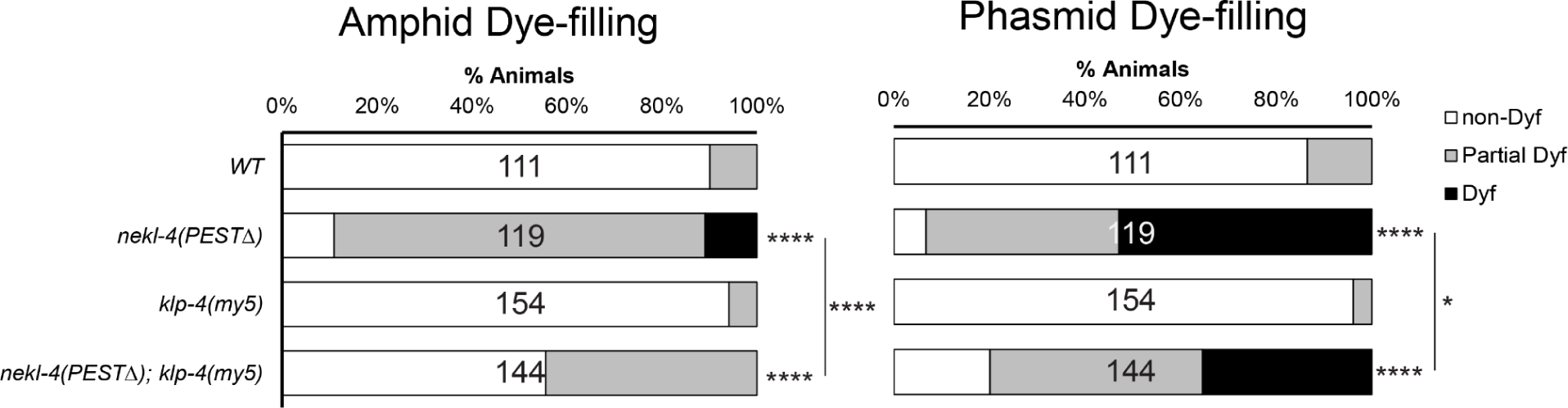
Mutation of kinesin-8 KLP-4/KIF13 suppresses the *nekl-4(PESTΔ)* Dyf phenotype. * indicates p ≤ 0.05, **** indicates p ≤ 0.0001 by Kruskal-Wallis one-way ANOVA with post hoc Dunn’s correction for multiple comparisons.

**Supplemental Figure 8.**
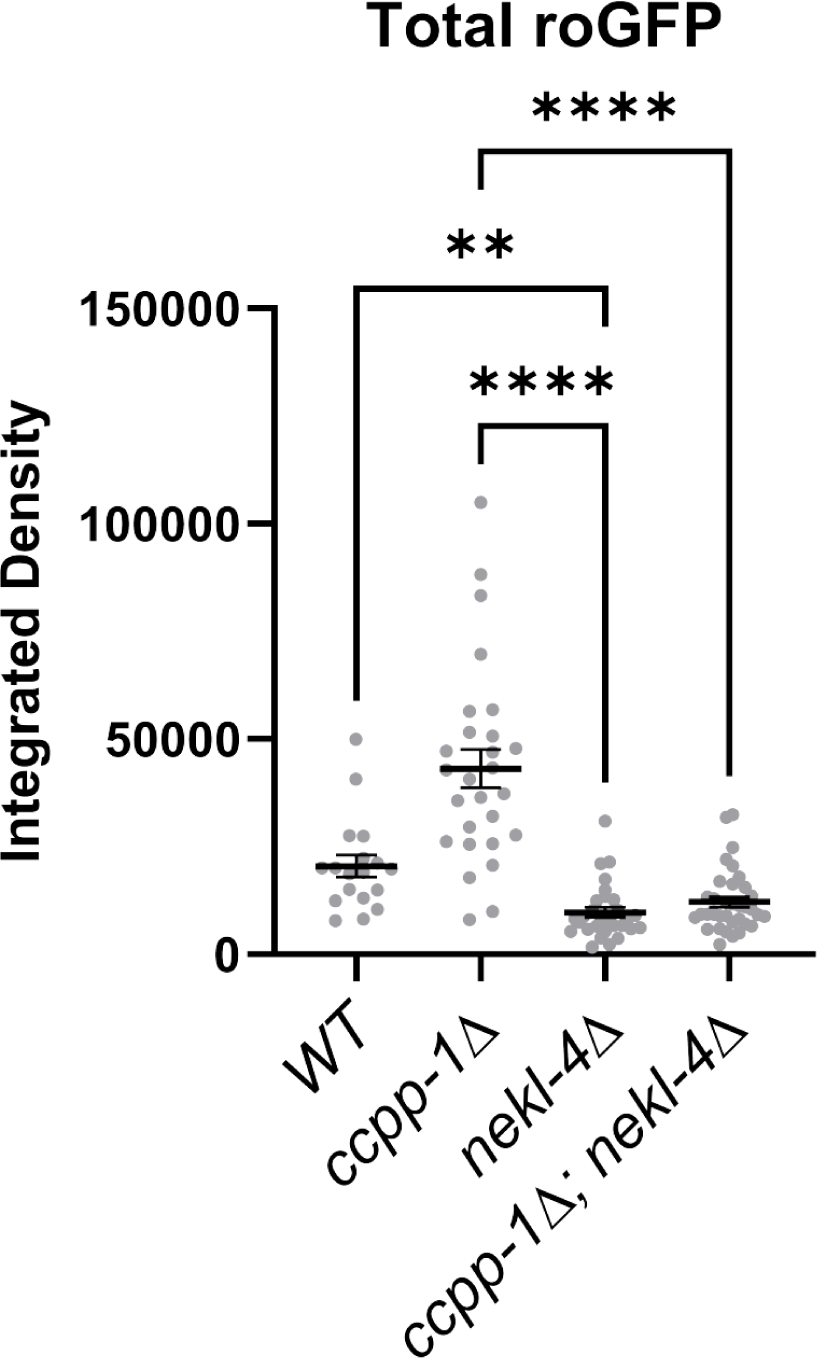
Total roGFP signal measured in phasmid soma. Calculated by adding the integrated densities of the 488 nm and 405 nm channels. Mean ± SEM; ** indicates p ≤ 0.01, **** indicates p ≤ 0.0001 by Kruskal-Wallis one-way ANOVA with post hoc Dunn’s correction for multiple comparisons.

**Supplemental Figure 9.**
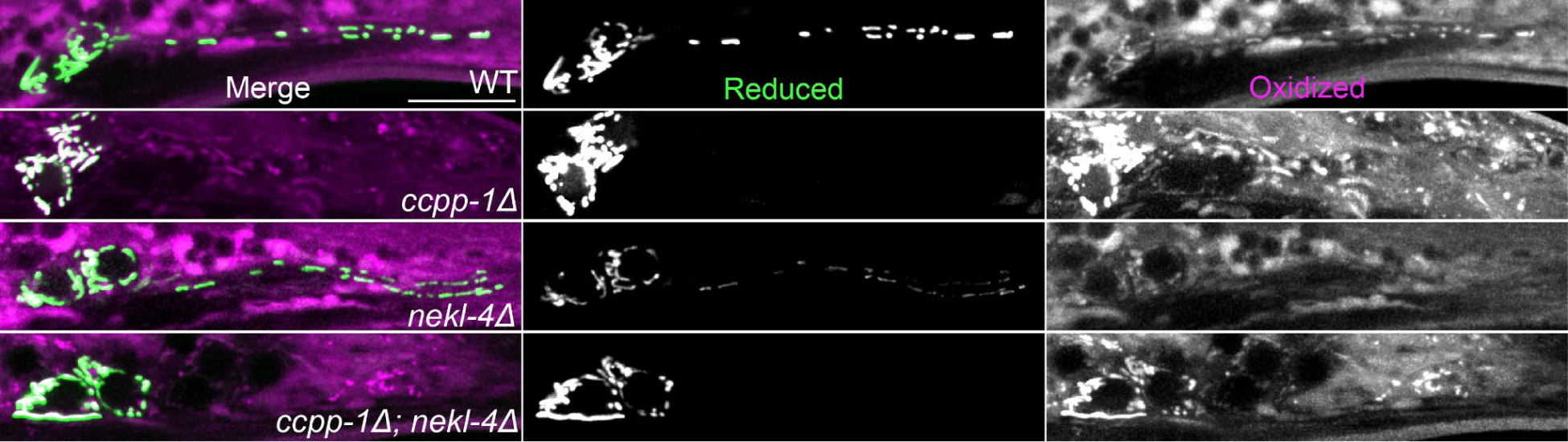
Mitochondrial transport is defective in *ccpp-1Δ* and *ccpp-1Δ; nekl-4Δ* mutants. Uncropped images of roGFP in the phasmid neurons. Scale bar = 10 µm.

## Acknowledgements

This work was supported by National Institutes of Health (NIH) DK059418, DK116606 and NS120745 (M.M.B), NIH NS122438 (K.M.P), and R24 OD010943 (D.H.H). We thank the members of the Barr and Rongo laboratories for valuable input on this research, particularly Inna Nikonorova, Juan Wang, Jonathan Dietz, Joelle Smart, and Tatiana Popovichenko. Additionally, K. M. P. thanks his thesis committee: Monica Driscoll, Marc Gartenberg, and Bonnie Firestein. We thank Gloria Androwski and Nanci Kane for technical assistance. Some strains were provided by the CGC, which is funded by NIH Office of Research Infrastructure Programs (P40 OD010440). We also thank WormBase (U41 HG002223) and WormAtlas (R24 OD010943) for online resources.

## Notes

### Competing Interest Statement

The authors have declared no competing interest.

